# Restoring O-glycosylation and expression of MUC2 limits progression of colorectal cancer

**DOI:** 10.1101/2024.01.25.577208

**Authors:** Yian Yang, Yuesong Yin, Wei Xu, Yan Kang, Jiawei Chen, Yanfeng Zou, Zhigang Xiao, Zheng Li, Peiguo Cao

**Affiliations:** Department of Oncology, the Third Xiangya Hospital, Central South University, Changsha, China; Department of Orthopedics, the Third Xiangya Hospital, Central South University, Changsha, China; NHC Key Laboratory of Carcinogenesis, Cancer Research Institute, School of Basic Medical Science, Central South University, Changsha, China; Department of General Surgery, the First Affiliated Hospital of Hunan Normal University (Hunan Provincial People’s Hospital), Changsha, China

**Author notes:** Correspondence: Dr. Peiguo Cao,; Dr. Zheng Li.

**Keywords:** Colorectal Cancer, MUC2, O-glycosylation, B3GNT6, Rosiglitazone, Intestinal Barrier

## Abstract

This study investigated the molecular mechanisms underlying the regulation of MUC2 expression and O-glycosylation modification in colorectal cancer. In addition, the potential of rosiglitazone to inhibit colorectal cancer by improving MUC2 glycosylation to protect intestinal barrier function was explored. In vitro, lectin staining combined with Co-IP assay was used to detect glycosyltransferases regulating MUC2 O-glycosylation. ChIP and Luciferase experiments were used to verify the transcription factors regulating MUC2 expression level. Samples from CRC patients were used to detect differences in multimolecular expression. The AOM/DSS mouse model was used to validate the effect of rosiglitazone on inhibiting colorectal cancer progression. Our results showed that B3GNT6 acts as a glycosyltransferase to enhance the O-glycosylation level of MUC2 and maintain protein stability to resist degradation by StcE secreting from pathogenic bacteria. Furthermore, KLF4 directly promotes the transcription of B3GNT6 and MUC2, which are regulated by PPARg. Rosiglitazone activated PPARg-KLF4-B3GNT6 axis which increased the expression level and glycosylation of MUC2 and further improved the intestinal mucosal barrier function to delay the development of colorectal cancer in mice. These data suggest that O-glycosylation and expression of MUC2 is key to the maintenance of functional intestinal mucosa and rosiglitazone is a potential colorectal cancer therapeutic agent.

## Introduction

Colorectal cancer (CRC) is the fourth leading cause of cancer-related mortality worldwide^1^. The development of CRC is closely linked to a number of genetic and environmental factors, wherein an abnormal intestinal barrier is an important driver^2,3^. Intestinal dysfunction, inflammatory bowel disease, and celiac disease are all associated with abnormal intestinal barrier function, which leads to increased intestinal permeability, bacterial translocation, a disturbed internal environment, and the growth of adenomas, which may eventually contribute to colorectal cancer^4,5^.

MUC2 is a member of the mucin family and is processed MUC2 is densely packaged and stored in secretory granules or vesicles, transported to the cell surface, and released into the intestinal lumen. MUC2 combines with large amounts of water and other substances to form mucus gel, which is a critical part of the intestinal barrier^6,7^. Reduced expression of MUC2 in the intestinal lumen can lead to increased permeability of the intestinal mucosa and impairment of the intestinal barrier^8–10^.

The function of MUC2 is influenced by glycosylation, the most important post-translational modification^11,12^. Highly glycosylated modifications ensure that MUC2 is able to form gels when combined with substantial quantities of water, thus playing a critical role in the production of intestinal mucus. It has been shown that aberrant glycosylation of the MUC2 protein is responsible for the inability to produce a normal mucus structure^13,14^. However, neither the abnormal changes in the glycosylation modifications of MUC2 in colorectal carcinogenesis nor the underlying mechanisms have been investigated.

In this study, we found that the transcriptional expression and O-glycosylation level of MUC2 protein were regulated by the cascade of KLF4 and B3GNT6. In addition, the PPARg agonist rosiglitazone could restore intestinal barrier function and inhibit the progression of colorectal cancer by acting MUC2 via KLF4-B3GNT6 axis. These findings suggest a potential novel treatment for colorectal cancer patients through the recovery of intestinal barrier functions.

## Results

### 1. MUC2 expression and O-glycosylation at a low level in colorectal cancer

We analyzed the mRNA expression and distribution of MUC2 across organs in the human body using GEPIA2 and found that MUC2 is mainly expressed in the digestive tract (Figure EV 1.A). RNA-seq data and IHC results of MUC2 from colorectal cancer patients in the TCGA database (https://portal.gdc.cancer.gov) showed significantly low expression of MUC2 in colorectal cancer tissues (Figure EV 1.B-C). Patients with low MUC2 expression had a poor prognosis in terms of overall survival (Figure EV 1.D). We collected 31 pairs of colorectal cancer patients’ tumors and adjacent tissues for Alcian blue staining (Figure 1.A-D) and Western Blot (Figure 1.E-F). The results revealed that MUC2 expression levels were significantly reduced in the tumor tissues compared with the adjacent tissues with the thinning of the mucus layer.

**Figure 1.**
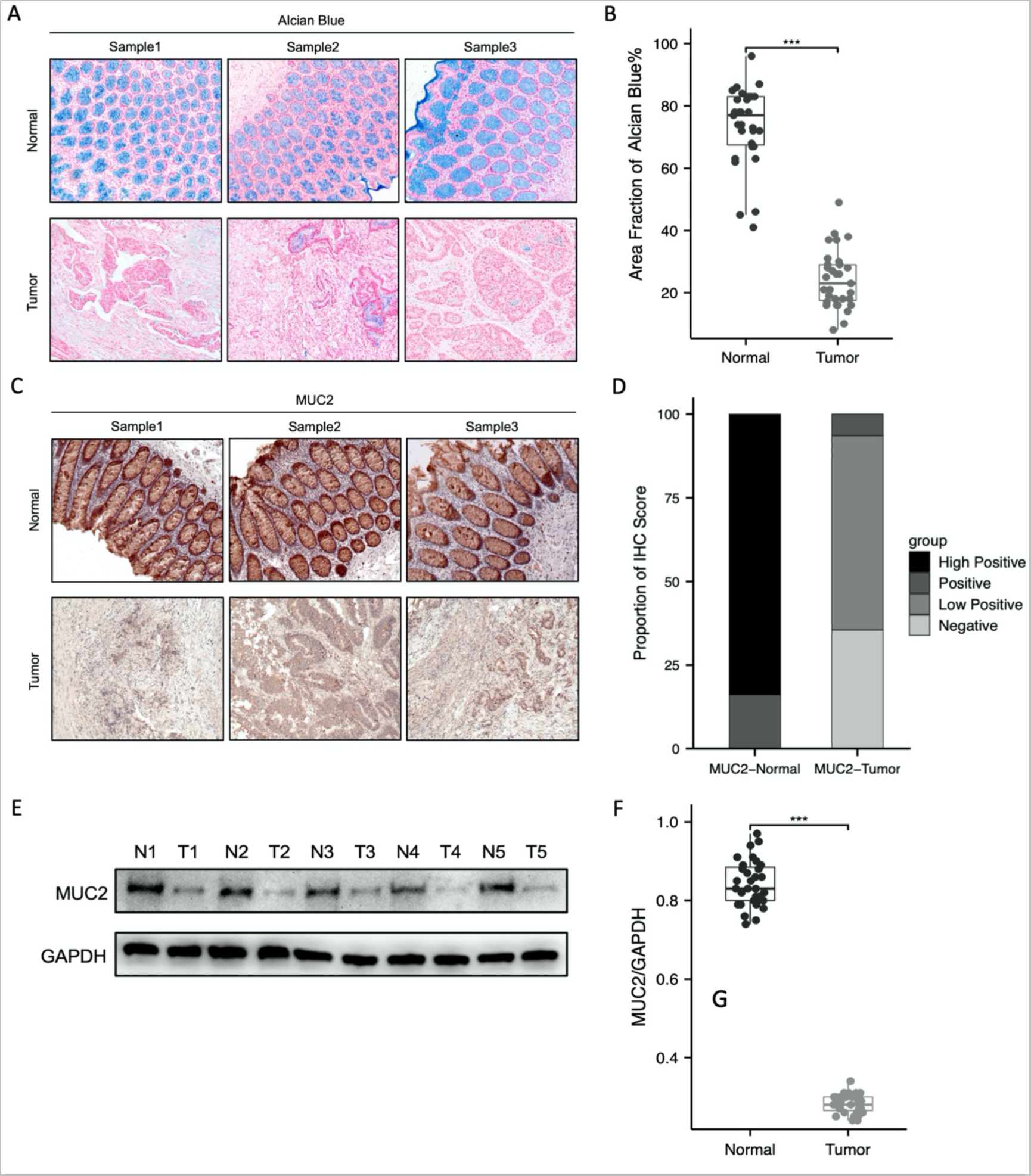
The expression of MUC2 is low in colorectal cancer. (A-B) Detection and quantification of intestinal mucus secretion by Alcian Blue staining (n=31, normal tissue; n=31, tumor tissue). Scale bars, 50 µm. (C-D) IHC staining and quantitation of MUC2 expression in the CRC patients (n=31, normal tissue; n=31, tumor tissue). Scale bars, 50 µm. (E) Western Blot analysis of MUC2 and (F) band densitometry quantifications in the CRC patients (n=31, normal tissue; n=31, tumor tissue). Data were analyzed by ordinary one-way ANOVA with Tukey’s multiple comparisons. The error bars indicate means ±standard error (SEM). *p<0.05, **p<0.01, ***p<0.001. ANOVA, analysis of variance; IHC, immunohistochemistry; CRC, Colorectal Cancer.

The further detection found that MUC2 expression levels were significantly lower in colorectal cancer cell lines than in normal cell lines (Figure EV 1.E). Since MUC2 is a large protein (5179 amino acids and a molecular mass of 540 kDa), it is difficult to construct a plasmid containing the full length of the MUC2 transcript. We used siRNA to reduce the expression of MUC2 to verify its effect on colorectal cancer cells (Figure EV 1.F). The observations of clonogenicity and CCK8 proliferation assays suggested that MUC2 silencing resulted in an increase in the number of clone-forming colonies and cell proliferation capacity (Figure EV 1.G-H). Together, these data demonstrated that MUC2 is downregulated in colorectal cancer cells and inhibition of MUC2 can promote cell proliferation.

### 2. B3GNT6 regulates MUC2 O-glycosylation modifications and protects its protein stability

Pearson statistical analysis of RNA-seq data from TCGA-COAD was used to identify genes associated with MUC2 (p<0.05), and the top 100 positively associated genes of MUC2 were obtained based on correlation coefficients (Figure EV 2.A-B). KEGG analysis suggested that the function of MUC2-associated genes was enriched in the O-glycosylation pathway (Figure EV 2.C). Recent studies has found that aberrant O-glycosylation is associated with colorectal carcinogenesis and decreased expression of MUC2^18^. However, the underlying mechanisms of O-glycosylation and MUC2 in CRC have not been investigated. The results of GSEA and PPI network interaction analysis also showed that MUC2 was closely related to a series of glycosyltransferases, and B3GNT6 ranked first (Figure 2.A-B). RNA-seq data from TCGA-COAD were analyzed for downregulation of B3GNT6 in colorectal cancer tissues and a positive correlation with MUC2 expression levels (correlation coefficient=0.813, p<0.001) (Figure EV 3.A-B). The results of the IHC of the above 31 paired CRC samples showed that B3GNT6 expression was significantly reduced or even absent in tumor samples (Figure 2.C-D). The O-glycosylation of mucin can be further divided into four main core structures (Core 1-4), which have different tissue specificity and physiological functions (Figure EV 3.C). B3GNT6 is responsible for creating the core 3 structure of O-glycans, which are important components of mucin-type glycoproteins^19^. We hypothesized that B3GNT6 could modulate the O-glycosylation modification of MUC2 and influence its expression in colorectal cancer tissues.

**Figure 2.**
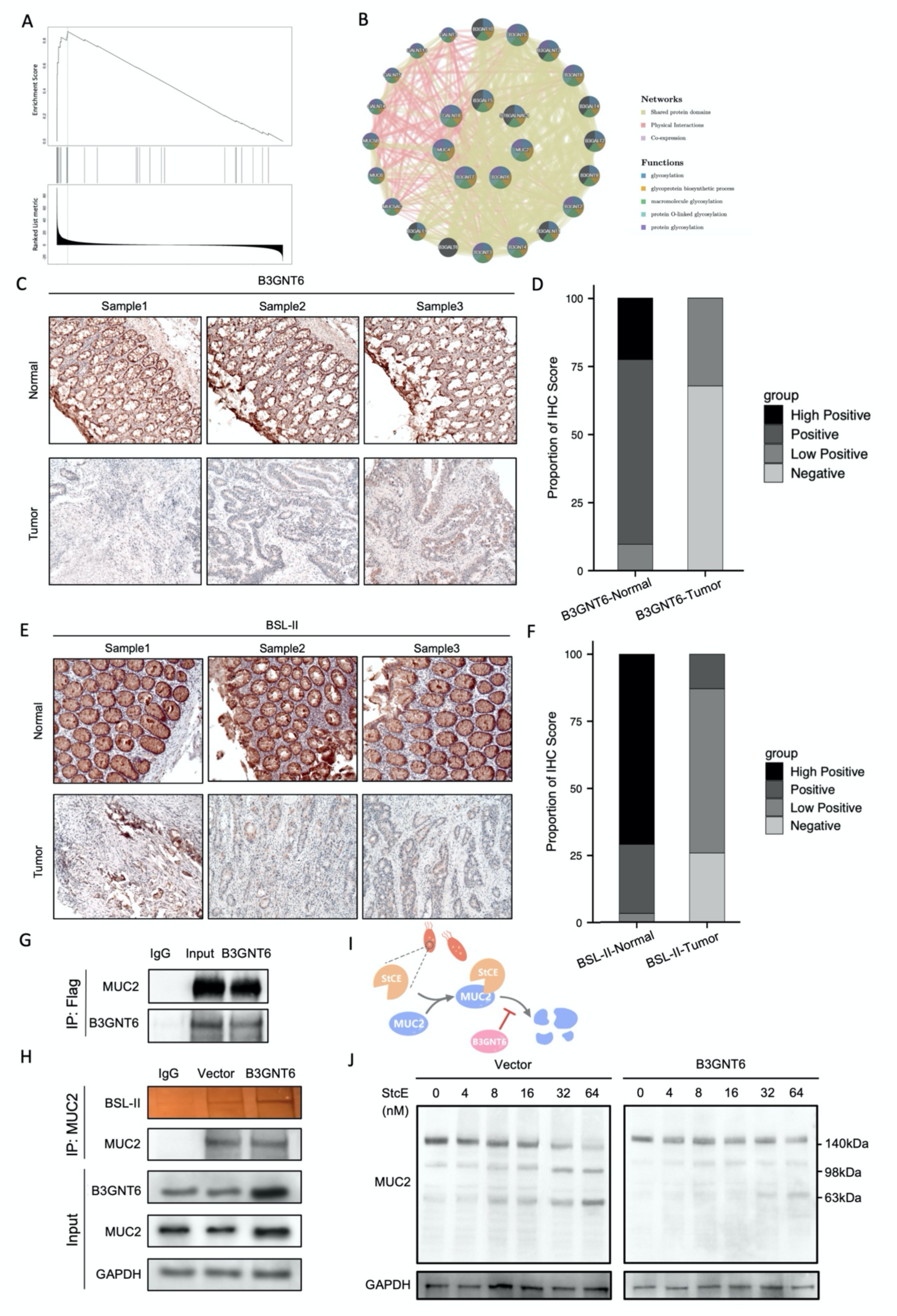
B3GNT6 enhances the expression and stability of MUC2 protein. (A) GSEA was performed to analyze a specific enrichment of factors in the Mucin type O-glycan biosynthesis pathway. (B) PPI network was analyzed with GeneMANIA. (C-D) IHC staining and quantitation of B3GNT6 in the CRC patients. Scale bars, 50 µm. (E-F) Lectin staining using BSL-II and quantitation of Core3 in the CRC. (G) Co-IP analysis to detect interaction of MUC2 and B3GNT6 after transfection of pcDNA3.1-B3GNT6-Flag vector, the protein complex bound to B3GNT6 was pulled down by anti-flag antibody (n=3, Input; n=3, IgG; n=3, B3GNT6). (H) Co-IP and lectin staining analysis to detect the expression of Core3 in MUC2 after transfection of pcDNA3.1-B3GNT6-Flag vector, the protein complex was pulled down by MUC2 antibody (n=3, Input; n=3, IgG; n=3, B3GNT6). (I) StcE specifically cleaves MUC2 O-glycosylation modified peptides, resulting in MUC2 degradation. (J) Western Blot to detect the degradation of MUC2 after the LOVO cells were treated with different concentrations of StcE (0nM, 4nM, 8nM, 16nM, 32nM, 64nM) for 3 hours. Data were analyzed by ordinary one-way ANOVA with Tukey’s multiple comparisons. The error bars indicate means ±standard error (SEM). *p<0.05, **p<0.01, ***p<0.001. ANOVA, analysis of variance; PPI, the protein–protein interaction; GSEA, gene set enrichment analysis, IHC, immunohistochemistry; Co-IP, co-immunoprecipitation; StcE, Recombinant Escherichia coli O157: H7 Metalloprotease.

The mRNA and protein levels of MUC2 were increased in LOVO cells following in vitro construction of the pcDNA3.1-B3GNT6 vector to overexpress B3GNT6 (Figure EV 3.D-E). Bandeiraea (Griffonia) Simplicifolia Lectin II (BSL-II) can recognize α- or β-linked N-acetylglucosamine residues on the nonreducing terminal of oligosaccharides and specifically binds Core3 O-glycan^20^. Through biotin-labeled BSL-II detection, we found the overall level of Core3 O-glycosylation modification was lower in colorectal cancer patients than in adjacent tissues (Figure 2.E-F). The same lectin staining of LOVO cells overexpressing B3GNT6 revealed a significant increase in Core3 modification (Figure EV 4.A). Furthermore, immunoprecipitation experiments on LOVO cells transfected with the pcDNA3.1-B3GNT6 vector showed that B3GNT6 can directly interact with MUC2 (Figure 2.G). The BSL-II lectin staining demonstrated that overexpression of B3GNT6 caused a significant increase in the level of Core3 O-glycosylation in MUC2 (Figure 2.H).

StcE is secreted by a pathogenic bacterium, enterohaemorrhagic Escherichia coli (EHEC). In the process of intestinal infection with EHEC, StcE can promote the adhesion of EHEC to the surface of epithelial cells and inhibit neutrophil adhesion to epithelial cells^21^. StcE recognizes S/T-X-S/T polypeptide sequences containing densely modified O-glycosylation regions and cleaves the peptide when O-glycosylation occurs at the first S/T site in the peptide sequence^22,23^. Interestingly, small degraded fragments of MUC2 induced by StcE were significantly decreased after overexpression of B3GNT6 in LOVO cells (Figure 2.I-J). Meanwhile, ELISA analysis showed that the concentration of MUC2 in the media was also enhanced (Figure EV 4.B-C). These results suggested that B3GNT6 mediated Core3 O-glycosylation of MUC2 and kept it stable in StcE-induced degradation, which may restore the intestinal barrier function of patients with colorectal cancer.

### 3. MUC2 and B3GNT6 is regulated by KLF4

To further investigate the mechanism of low MUC2 expression in colorectal cancer, we analyzed the Jaspar database (http://jaspar.genereg.net) to predict the transcription factors that could specifically bind to the MUC2 promoter. Among which the Krüppel-like factor (KLF4) as a tumor suppressor gene in colorectal cancer was positively correlated with MUC2 expression (Figure EV 5.A-B). After transfecting KLF4-specific siRNA or pcDNA3.1-KLF4 plasmid into LOVO cells, the mRNA and protein levels of MUC2 was altered in accordance with KLF4 changes compared to the control group (Figure EV 5.C-F). Western blot assays also showed that the protein expression levels of MUC2 gradually decreased with increasing concentration of the KLF4 inhibitor Kenpaullone (Figure 3.A). Using the above 31 paired CRC sample cohort, we found that KLF4 was downregulated in tumor tissues compared to adjacent tissues (Figure 3.B-C). ChIP-qPCR and PCR assays validated the candidate binding target site (76-87 bp upstream of the ATG initiation codon of MUC2) (Figure EV 5.G) (Figure 3.D-E). To further confirm the KLF4 binding site in the MUC2 promoter region, the promoter region of MUC2 (-2,000 to +100 bp) was cloned into the pGL3-Basic luciferase reporter plasmid (WT) and the binding site was mutated (Mut) (Figure EV 5.H). Luciferase reporter assay demonstrated that the mutation completely eliminated luciferin (Figure 3.F).

**Figure 3.**
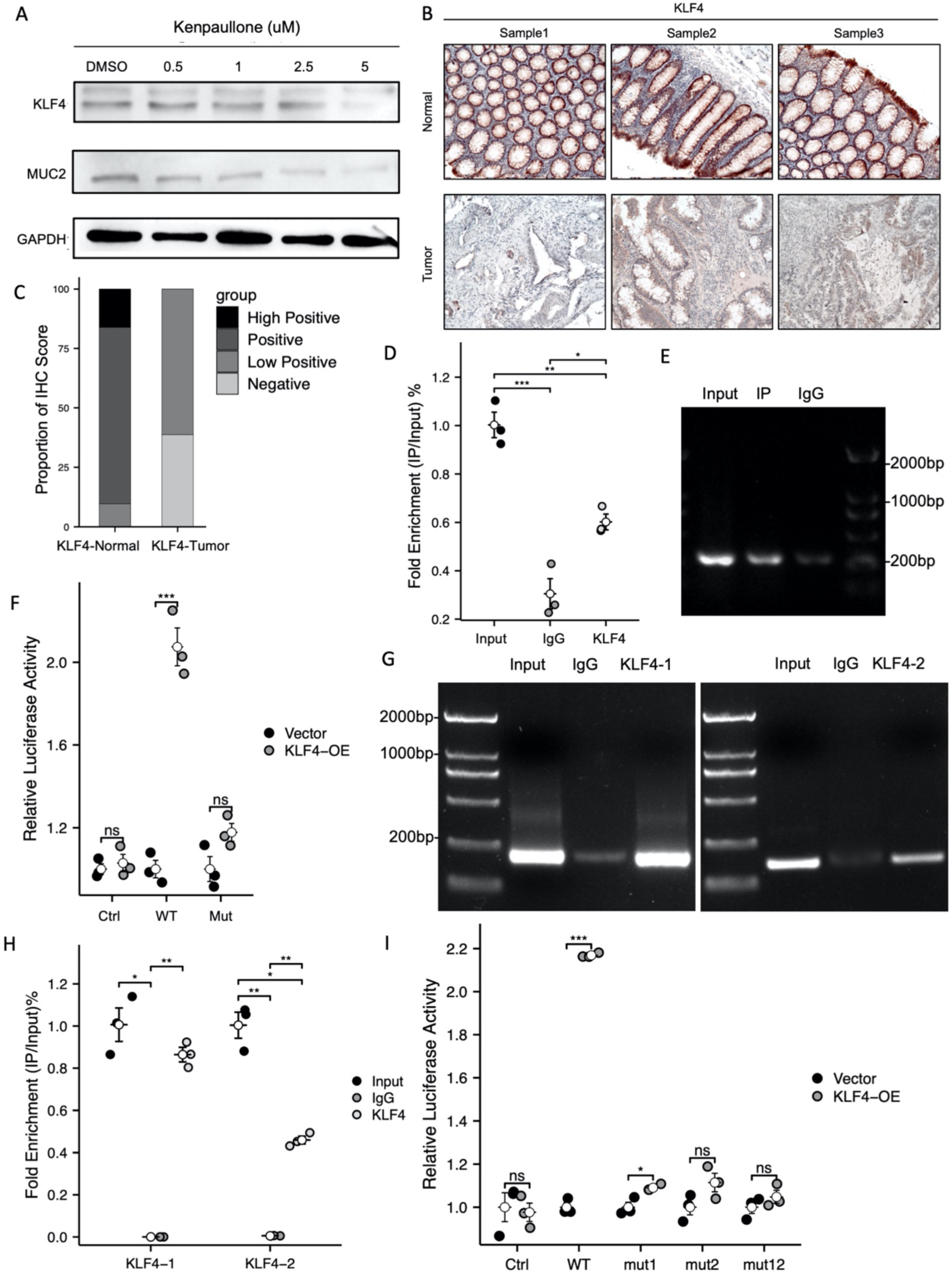
KLF4 directly binds to the MUC2 and B3GNT6 promoter to promote its transcription. (A) Western Blot to detect the expression of MUC2 after the cells were treated with different concentrations of KLF4 inhibitor Kenpaullone (0uM, 0.5uM, 1uM, 2.5uM, 5uM) for 24 hours. (B-C) IHC staining and quantitation of KLF4 in the CRC patients (n=31, normal tissue; n=31, tumor tissue). Scale bars, 50 µm. (D-E) ChIP enrichment was quantified using qPCR and PCR analysis to detect MUC2-binding site (n=3, Input; n=3, IgG; n=3, CHIP-KLF4). (F) The above plasmids and pCMV-Renilla plasmid were co-transfected into LOVO cells, and the normalized luciferase activity was measured (n=5, Ctrl; n=5, WT; n=5, mut). (G-H) ChIP enrichment was quantified using qPCR and PCR analysis to detect B3GNT6-binding sites (n=3, Input; n=3, IgG; n=3, CHIP-KLF4). (I) The above plasmids and pCMV-Renilla plasmid were co-transfected into LOVO cells, and the normalized luciferase activity was measured (n=5, Ctrl; n=5, WT; n=5, mut1; n=5, mut2; n=5, mut12).Data were analyzed by ordinary one-way ANOVA with Tukey’s multiple comparisons. The error bars indicate means ±standard error (SEM). *p<0.05, **p<0.01, ***p<0.001. ANOVA, analysis of variance; qPCR, quantitative polymerase chain reaction; ChIP, chromatin immunoprecipitation; CRC, Colorectal Cancer.

Interestingly, we also found a significant positive correlation between B3GNT6 and KLF4 in the TCGA-COAD dataset (Figure EV 6.A). Overexpression of KLF4 could promote B3GNT6 expression, whereas knockdown of KLF4 could inhibit B3GNT6 expression in LOVO cells (Figure EV 6.B-C). Furthermore, KLF4 binding motifs were predicted that there were two binding sites in front of the B3GNT6 ATG initiation codon (1924-1934 bp and 155-165 bp) (Figure EV 6.D), ChIP assay also verified this prediction (Figure 3.G-H). The luciferase reporter plasmid (WT) containing the B3GNT6 promoter region was designed, and two candidate binding targets (mut1, mut2 and mut12) were mutated separately or simultaneously (Figure EV 6.E). The single and double mutants with two binding sites significantly decreased the luciferase activity compared to the wild-type control (Figure 3.I). These above-mentioned data suggested that KLF4 can stimulate the transcription of B3GNT6 while directly activating the transcription of MUC2.

### 5. PPARg activates MUC2 and B3GNT6 expression through KLF4

We want to find drugs targeting MUC2 to restore the intestinal barrier function in CRC patients. However, there are no known drugs that directly target MUC2, B3GNT6, or KLF4. It has been demonstrated that the transcriptional regulator PPARg can regulate the expression of KLF4 in pancreatic cancer cells^24,25^. TCGA-COAD data analysis showed that the expression of PPARg was low in colorectal cancer, and there was a significant positive correlation between PPARg and KLF4 in CRC samples (Figure EV 7.A-C). IHC analysis confirmed the reduced expression of PPARg in tumor tissues of CRC patients (Figure 4.A-B). Furthermore, we found that changes in the expression levels of KLF4, B3GNT6, and MUC2 were consistent with those of PPARg following treatment of LOVO cells with graded concentrations of the PPARg agonist rosiglitazone or the inhibitor GW9662 (Figure 4.C-D).

**Figure 4.**
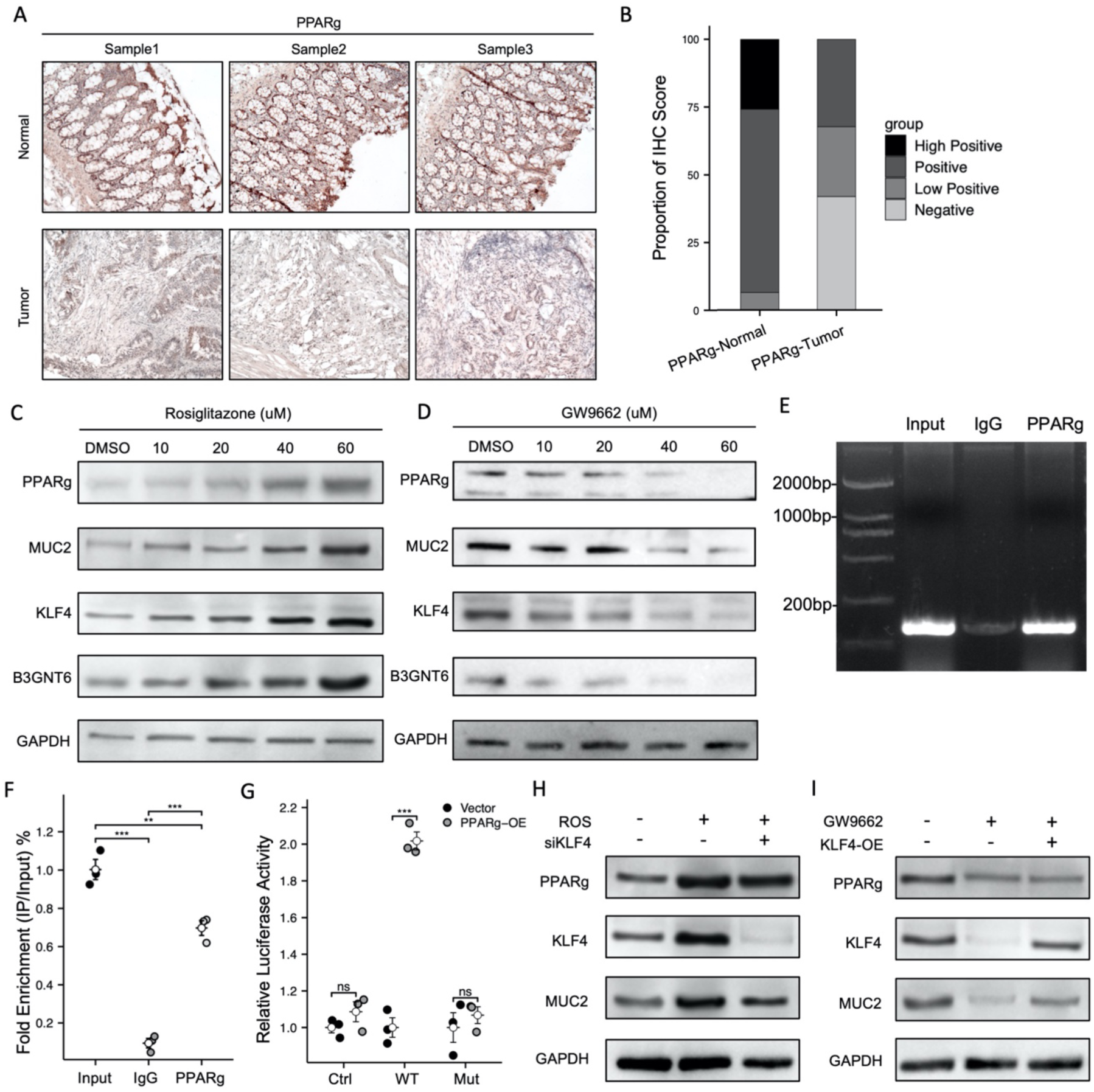
PPARg promotes the expression of MUC2 and B3GNT6 through KLF4. (A-B) IHC staining and quantitation of PPARg in the CRC patients (n=31, normal tissue; n=31, tumor tissue). Scale bars, 50 µm. (C-D) Western Blot to detect the expression levels of PPARg, MUC2, KLF4 and B3GNT6 after the cells were treated with different concentrations of Rosiglitazone (0, 10uM, 20uM, 40uM, 60uM) or GW9662 (0, 5uM, 10uM, 15uM, 20uM) for 24 hours. (E-F) ChIP enrichment was quantified using PCR and qPCR analysis to detect KLF4-binding site (n=3, Input; n=3, IgG; n=3, CHIP-PPARg). (G) The above plasmids and pCMV-Renilla plasmid were co-transfected into LOVO cells, and the normalized luciferase activity was measured (n=5, Ctrl; n=5, WT; n=5, mut). (H-I) Western Blot detect the expression levels of PPARg, KLF4 and MUC2 after the cells were treated with siRNA-KLF4 transfection and Rosiglitazone (40uM), or with pcDNA3.1-KLF4 plasmid transfection and GW9662 (20uM). Data were analyzed by ordinary one-way ANOVA with Tukey’s multiple comparisons. The error bars indicate means ±standard error (SEM). *p<0.05, **p<0.01, ***p<0.001. ANOVA, analysis of variance; IHC, immunohistochemistry; ChIP, chromatin immunoprecipitation; qPCR, quantitative polymerase chain reaction; CRC, Colorectal Cancer.

There were two candidate binding sites of PPARg in the MUC2 promoter region predicted by the Jaspar database (1959-1969 bp or 517-535 bp upstream of the ATG initiation codon). However, the ChIP experiment results suggested that PPARg could not directly bind with MUC2 promoter DNA (Figure EV 7.D-E). We therefore hypothesized that PPARg indirectly regulates the expression level of MUC2 through KLF4. PCR and qPCR experiments following ChIP assay demonstrated that PPARg directly regulates KLF4 transcription (Figure 4.E-F). Mutation of the PPARg binding site in the KLF4 promoter region of the luciferase reporter plasmid completely abolished luciferase activity (Figure EV 7.G) (Figure 4.G). In addition, we transfected siKLF4 into rosiglitazone-treated LOVO cells. We found the upregulation of MUC2 induced by rosiglitazone was decreased while KLF4 expression was inhibited (Figure 4.H). Similarly, when GW9662 was used to inhibit PPARg while KLF4 was persistently expressed, the expression level of MUC2 was higher than that of GW9662 alone (Figure 4.I). The above data suggested that PPARg regulates MUC2 expression via activating transcription of KLF4.

In vivo, diets containing rosiglitazone (10 mg/kg) or GW9662 (10 mg/kg) were used to feed C57BL/6 mice. Two weeks later, IHC assays revealed that PPARg, KLF4, B3GNT6, and MUC2 were all increased in the colon of mice in the rosiglitazone group, and the thickness of the mucus layer increased. We also observed that BSL-II lectin staining increased in the rosiglitazone group while being downregulated in the GW9662 group (Figure EV 8.A). Further collection of the intestinal mucus for ELISA analysis revealed a significant increase in MUC2 secretion in the rosiglitazone group of mice and a decrease in the GW9662 group (Figure EV 8.B). These results suggested that rosiglitazone active PPARg-KLF4-B3GNT6/MUC2 axis to increase MUC2 expression and O-glycosylation modification in vivo.

### 6. Rosiglitazone inhibits colorectal cancer development through the PPARg-KLF4-B3GNT6/MUC2 axis

Given the importance of rosiglitazone’s activation of the above-mentioned potential tumor suppression genes via agonistic PPARg expression, we assumed that rosiglitazone may be a therapeutic agent against colorectal cancer. To verify this hypothesis, the proliferation of CRC cells after administration of GW9662 or rosiglitazone was first measured in vitro using CCK8. The result was consistent with the prediction that GW9662 promoted CRC cell proliferation while rosiglitazone inhibited proliferation (Figure EV 9.A-B). With the further addition of pcDNA3.1-KLF4 plasmid to the GW9662 group or the addition of KLF4 siRNA to the rosiglitazone group, CCK8 assay showed that the effect of both drugs on the proliferation ability of colorectal cancer cells was eliminated (Figure EV 9.C-D). Therefore, we concluded that rosiglitazone inhibited the proliferation of colorectal cancer cells through KLF4.

To investigate the effect of rosiglitazone on the development of colorectal cancer in vivo, the AOM/DSS model of C57BL/6 mice was fed with rosiglitazone-containing fodder or ordinary fodder as a control. The mice in each group were continuously monitored for changes in body weight, and the rosiglitazone group showed a significant increase in body weight compared to the control group (Figure 5.A). According to the percentage of weight loss, fecal viscosity, and fecal occult blood, the disease activity index score (Disease Activity Index, DAI) was calculated. It was found that DAI in the rosiglitazone group was significantly lower than that in the control group after different DSS cycles (Figure 5.B). Meanwhile, the colon length was greater in the rosiglitazone group than the control group at the end of each DSS cycle (Figure 5.C-D). The inner surface of the colon was exposed by longitudinal sectioning and photographed, and the rosiglitazone group had significantly less tumor infiltration than the control group (Figure 5.E). The number of tumors in the colon of the mice was counted and found to be significantly lower in the rosiglitazone group than in the control group (Figure 5.F). At the same time, the tumor was divided into three grades according to its diameter (< 2mm, 2-4mm, > 4mm). The results showed that the number of tumors in the rosiglitazone group was lower than that in the control group (Figure 5.G). The tumorgenic mouse colons were sectioned in paraffin (Figure 5.H), and the pathological grade by HE staining was shown in the Table EV1. The results suggested that the histology score of the rosiglitazone group was lower than that of the control group at different DSS cycles (Figure 5.I). In conclusion, rosiglitazone could delay the progression of colorectal cancer in the AOM/DSS model and reduce tumor burden.

**Figure 5.**
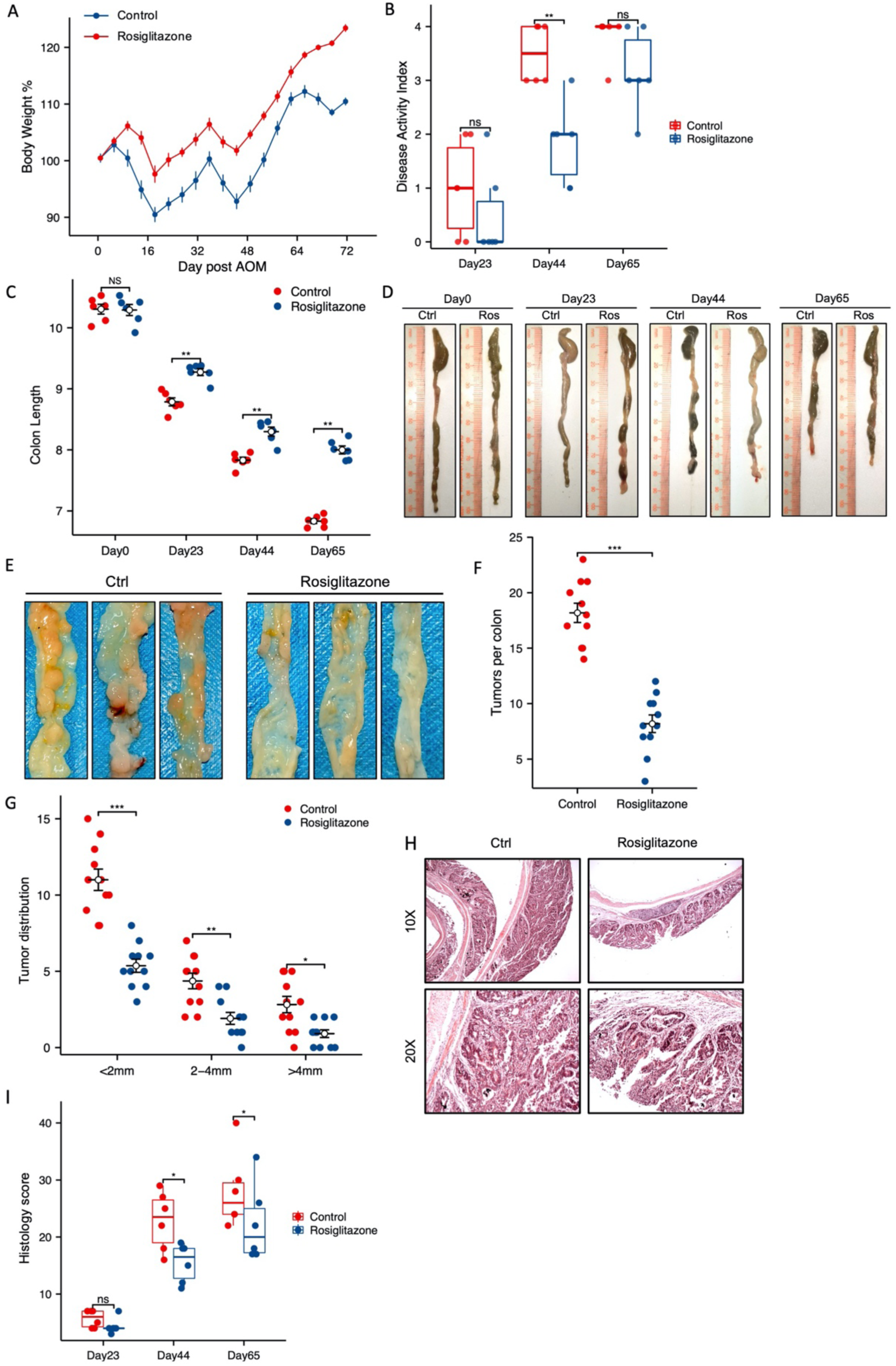
Rosiglitazone can inhibit the occurrence and development of colorectal cancer. (A) Line chart of standardized weight changes from the day of injection of AOM (n=6, AOM/DSS-Control; n=6, AOM/DSS-Rosiglitazone). (B) DAI score of mice (Day23, Day44, Day65 post AOM) according to the percentage of weight loss, fecal viscosity and fecal occult blood. (C) Colon length on day7, Day23, Day44, Day65 after AOM treatment. (D-E) Representative photographs of the colon of mice at the end of different periods (Day0, Day23, Day44, Day65) of DSS cycle and the colon of the colon of mice after AOM/DSS treatment. (F) Tumor number per colon in AOM/DSS treated mice. (G) Tumor distribution measured by the number of tumors in different diameter tumor groups (< 2mm, 2-4mm, > 4mm). (H) H&E staining and (I) Representative histological images of colon tissues (n=6, AOM/DSS-Ctrl; n=6, AOM/DSS-Rosiglitazone). Scale bars, 200 µm. Data were analyzed by ordinary one-way ANOVA with Tukey’s multiple comparisons. The error bars indicate means ±standard error (SEM). *p<0.05, **p<0.01, ***p<0.001. ANOVA, analysis of variance; DAI, Disease Activity Index; H&E, hematoxylin-eosin.

Colon tissues were dissected, and paraffin sections were prepared to detect changes in crypt structure and mucus secretion by HE staining and Alcian blue staining (Figure 6.A-B). The results showed that the rosiglitazone group was able to maintain normal crypt structure, and mucus secretion from the crypt persisted after the end of the first DSS cycle compared to the control group, suggesting that rosiglitazone delayed the loss of intestinal mucus barrier function during the pre-tumorigenic phase. IHC staining of colon sections from mice in the progressive stage showed that the expression levels of PPARg, KLF4, B3GNT6, MUC2, and Core3 modification in the control group gradually decreased or even disappeared, but the expression levels of these genes and Core3 modification in the rosiglitazone group maintain stability and some even increased before tumor formation (Figure 6.C-D). Ki67 staining results also showed the ability of rosiglitazone to significantly inhibit tumor proliferation (Figure 6.E-F). Together, rosiglitazone could maintain the intestinal mucosal barrier and prevent the development of colorectal cancer in the AOM/DSS model through regulation of PPARg-KLF4-B3GNT6/MUC2 axis.

**Figure 6.**
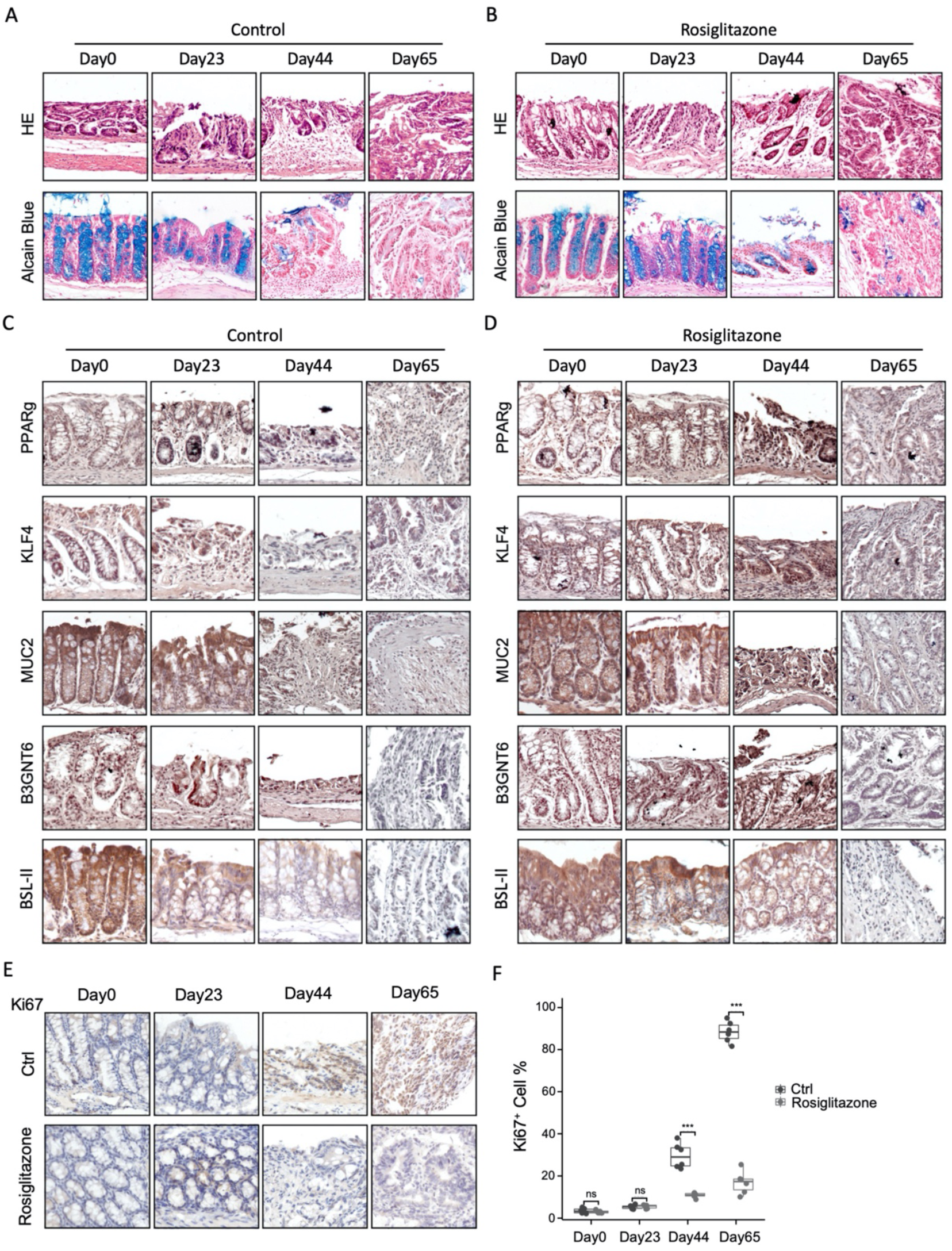
Rosiglitazone can reverse the sustained decrease in the expression of PAPRg, KLF4, MUC2 and B3GNT6 during the development of colorectal cancer. (A-D) Detection and quantification of intestinal mucus secretion by H&E staining, Alcian Blue staining and IHC staining in Intestinal tissues of mice at the end of different periods (Day0, Day23, Day44, Day65) of DSS cycle (n=6, AOM/DSS-Control; n=6, AOM/DSS-Rosiglitazone). Scale bars, 50 µm. (E-F) IHC-Ki67 staining and quantitation of the number of Ki67 positive cells in Intestinal tissues of mice at the end of different periods (Day0, Day23, Day44, Day65) of DSS cycle (n=6, AOM/DSS-Control; n=6, AOM/DSS-Rosiglitazone). Scale bars, 50 µm. Data were analyzed by ordinary one-way ANOVA with Tukey’s multiple comparisons. The error bars indicate means ±standard error (SEM). *p<0.05, **p<0.01, ***p<0.001. ANOVA, analysis of variance; H&E, hematoxylin-eosin; IHC, immunohistochemistry; CRC, Colorectal Cancer; ELISA, enzyme-linked immunosorbent assay.

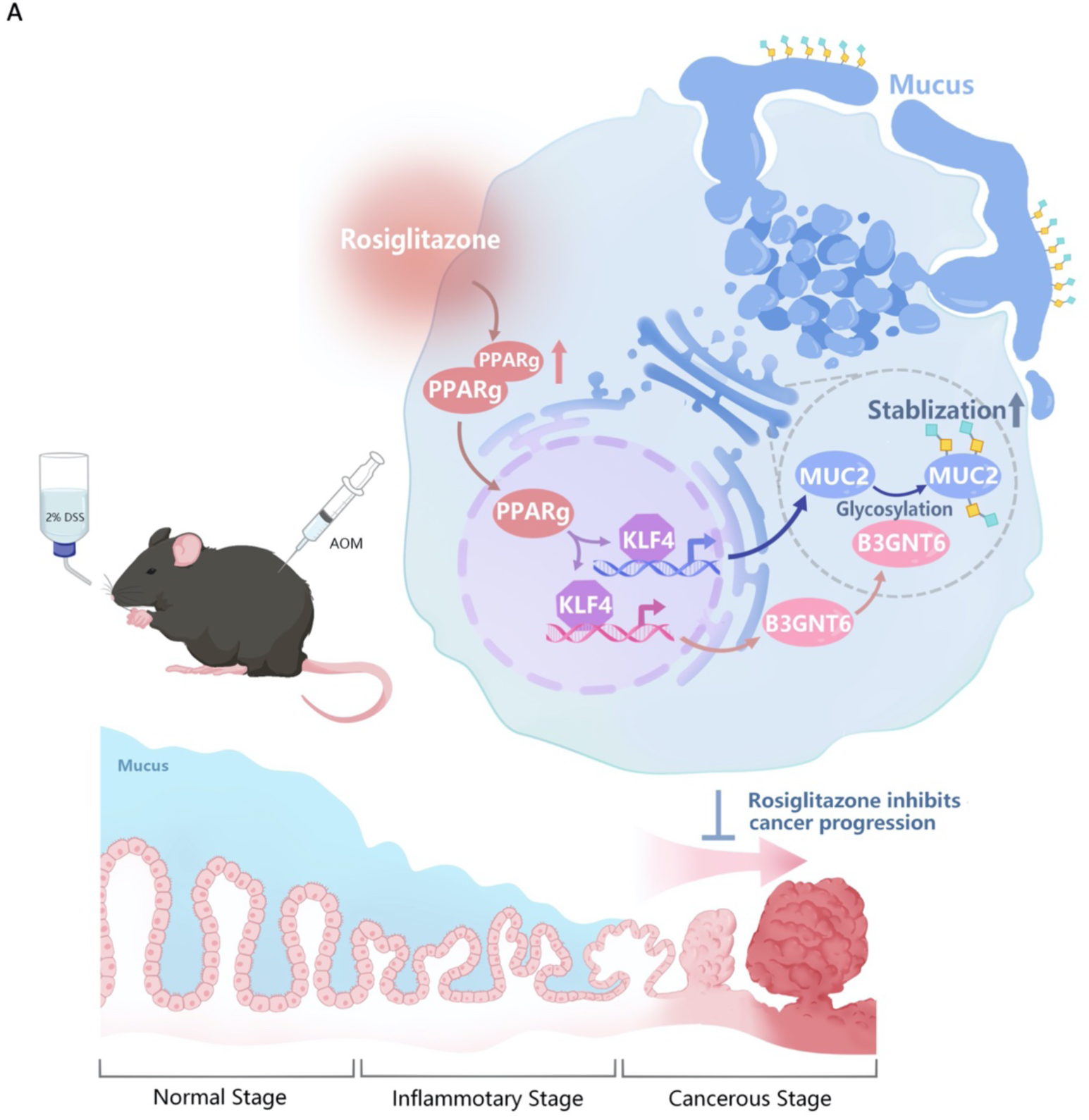

## Discussion

When the structure and function of the intestinal barrier are compromised, multiple alterations occur in the genome, metabolome, immunome, and flora, which in turn promote the development of colorectal cancer^26^. In this study, we have shown that KLF4 is responsible for promoting MUC2 transcription while also regulating the expression of B3GNT6 transcription to improve the O-glycosylation level of MUC2 protein as well as its protein stability, thereby maintaining the structural and functional integrity of the intestinal barrier. In the model of AOM/DSS-induced colon cancer, it was also shown that the expression levels of the above molecules decreased continuously with the disruption of the intestinal barrier and that rosiglitazone significantly slowed this process. Rosiglitazone may modulate the KLF4-B3GNT6/MUC2 axis through PPARg to inhibit colorectal cancer development.

Our study revealed a potential correlation between altered Core3 glycosylation levels of MUC2 in colorectal cancer and abnormal intestinal barrier function. B3GNT6, one of the essential glycosyltransferases, is responsible for the formation of mucopolysaccharides^27^. Our contribution is to elucidate the mechanism of action by which B3GNT6 interacts with MUC2 to increase the level of MUC2 O-glycosylation and improve MUC2 protein stability via resisting exogenous degradation from pathogenic bacterial secretion products. This is the novel aspect of our work.

Furthermore, by detecting colon tissue from patients and AOM/DSS model mice with the special Core3 modification staining agent BSL-II lectin, we found significantly reduced Core3 modification in cancerous tissue. This finding may be analogous to Sambucus nigra agglutinin (SNA) lectin^28^, Agaricus bisporus (ABA) lectin^29^, as well as other lectins as potential biomarkers^30^ for the early diagnosis of colorectal cancer. The B3GNT6-induced changes in Core3 glycosylation sites and MUC2 abundance can be further investigated in the future.

PPARg is a key gene in several pathways, including lipid differentiation, glucose metabolism homeostasis, and inflammatory response. It has been shown that activation of PPARg signaling can inhibit the proliferation of pathogenic flora and the development of colitis^31,32^. This innovative study investigated the molecular mechanism by which PPARg regulates the transcription of KLF4 to enhance B3GNT6 and MUC2 expression and maintain intestinal barrier homeostasis to impede colorectal cancer progression. KLF4 is responsible for regulating cell proliferation, apoptosis, differentiation, and tissue homeostasis in the body^33^. Reduced KLF4 expression has been shown to have a significant impact on both overall survival and recurrence rates in colorectal cancer patients^34^. The expression of genes involved in the cell cycle can be modulated by KLF4, which has been shown to suppress colorectal cancer progerssion^35,36^. Interestingly, PPARg is able to regulate KLF4 to simultaneously maintain the transcriptional level and protein glycosylation of MUC2. This is necessary for the stability of MUC2 and its efficient secretion into the intestinal lumen to maintain the function of the intestinal barrier. Therefore, the present study investigated the protective role of rosiglitazone activation of the PPARg-KLF4-B3GNT6/MUC2 signaling pathway in the progression of colitis to cancer.

Rosiglitazone is a thiazolidinedione with the potential to act as a high-affinity agonist of PPARg^37^. In an AOM/DSS-induced model of colorectal cancer, its administration resulted in a significant reduction in the rate of colorectal carcinogenesis and increased the thickness of intestinal mucosa in mice. Rosiglitazone is currently used as a treatment option for type II diabetes mellitus. A phase IV clinical trial showed that rosiglitazone had significant efficacy in the treatment of inflammatory bowel disease in 105 people who were in the mildly or moderately active stages of the disease^38^. Rosiglitazone is currently being tested in seven different clinical trials in people with various types of cancer, including breast, bladder, and prostate cancer. However, there are no reports of research into the use of PPARg agonists in the treatment of colorectal cancer. This is the first study to investigate the function and mechanism of rosiglitazone as a potential treatment for colorectal cancer.

The current treatment options for people with colorectal cancer who have an abnormal intestinal barrier are quite limited, and there are no successful anti-cancer drugs that activate the protection function provided by the intestinal barrier. This research reveals the KLF4-B3GNT6/MUC2 pathways and targeted drugs required to maintain intestinal barrier homeostasis, which has the potential to become a new diagnostic and therapeutic breakthrough for colorectal cancer.

## Materials and methods

### Statistics

Data values for each group were obtained from three or more independent measurements and expressed as mean±SD. all statistical analyses were performed using GraphPad Prism v.7.0. Statistical tests between groups were performed using Student’s t-test or ANNOVA. Variance was similar between group statistically compared. A p value of <0.05 was considered statistically significant, *p ≤ 0.05, **p ≤ 0.01, ***p ≤ 0.001.

### Study approval

The samples of patients were obtained and used with the approval of the Ethics Committee of the Third Xiangya Hospital of Central South University (ID: IRBS2023049). All animal models were constructed in accordance with the Regulations for the Administration of Affairs Concerning Experimental Animals established by State Scientific and Technological Commission of The People’s Republic of China and passed the animal ethical review by the Department of laboratory animals of Central South University (ID: CSU-2022-0684).

### External data set analysis

RNAseq data from the TCGA database (https://portal.gdc.cancer.gov) were downloaded and collated from the TCGA-COAD project and extracted in TPM format. Appropriate statistical methods were selected for statistical gene expression and visualized according to the data format characteristics. For survival analysis, clinical information of the TCGA-COAD project was obtained for Kaplan-Meier log Rank survival test. Spearman’s correlation analysis was applied in LinkedOmics (http://www.linkedomics.org/) to verify the correlations between MUC2 and identified target genes involved in significant pathways. GO term enrichment and KEGG pathway analysis of the top 100 genes in terms of correlation coefficient were performed with DAVID. PPI interaction networks were calculated analyzed by using GeneMANIA (http://genemania.org).

### Patient Samples

The colorectal samples were obtained from patients of the Department of Oncology at the Third Xiangya Hospital of Central South University, and included samples of tumor tissues and paracancerous tissue (greater than ≥5 cm from the tumor border) from 31 patients with colorectal cancer. The samples were obtained and used with the approval of the Ethics Committee of the Third Xiangya Hospital of Central South University (ID: IRBS2023049). All patients signed a written informed consent form and ensured the anonymity and confidentiality of their personal data. All patients had not received palliative surgery, chemotherapy or radiotherapy before radical surgery and all had been confirmed by postoperative pathological examination.

### Immunohistochemistry (IHC) assay

Colorectal tissues were fixed in 4% paraformaldehyde for 24 hours at room temperature and dehydrated by immersion in 70%, 85%, 95% and 100% ethanol for more than 30 minutes in sequence. The tissues were soaked in a 1:1 mixture of ethanol and xylene and xylene for more than 30 minutes to make them transparent. Paraffin sections were soaked in liquid paraffin wax for more than 30 minutes and then paraffin-embedded using an embedding mold, frozen at -20°C for 12 hours, and sectioned at a thickness of 4 um. Paraffin sections were baked in an oven at 65 °C for 30 minutes and re-hydrated by immersion in ethanol at 100%, 95%, 85%, 75%, and 0% concentrations for 2 minutes in that order. After re-hydrated, antigen retrieval was performed in sodium citrate buffer (Biosharp, BL619A) for 10 minutes and cooled to room temperature in PBS. Sections were soaked for 15 min using PBS containing 0.1% TritonX-100 followed by 5 washes of PBS for 4 min each. Endogenous Peroxidase Blocking Buffer (Beyotime, P0100B) was added dropwise to the sections and incubated for 10 min. After washing, Blocking Buffer for Immunol Staining (Beyotime, P0260) was added dropwise to the sections and incubated for 10 min. after washing, primary antibody was added dropwise and incubated overnight at 4°C. Antibody diluent (New Cell & Molecular Biotech, WB500D). After washing, the sections were incubated for 2 h at room temperature with biotin-labeled goat anti-rabbit IgG (Beyotime, A0279) at a dilution ratio of 1:250 with secondary antibody. Add dropwise to the sections and incubate for 1 h at room temperature. Prepare DAB working solution according to the instructions of DAB staining kit (ZSGB-BIO, ZLI-9018), add dropwise to the sections and incubate for 10 min at room temperature, then wash twice to terminate the incubation. Sections were dehydrated by immersion in 50%, 75%, 85%, 95%, 100%, xylene for 30 seconds, and immunohistochemistry was fixated by neutral balsam (Biosharp, BL704A) to cover the tissue. Immunohistochemical sections were imaged using a microscope image system (Zeiss, Axioscope 5). The staining in the IHC images was analyzed and scored by the IHC Profiler plugin in ImageJ software (Maryland, USA). Images were scored according to staining intensity versus staining area, High positive (3+), Positive (2+), Low Positive (1+) and Negative (0). The number of positive Ki67 stained cells was counted by the Tranable Weka Segmentation plugin.

### Alcian Blue staining

To show the structure of mucus secreted by cupped cells in colonic tissue, AB staining was performed according to the protocol of the Alcian Blue staining kit (Bioss, S0134), and stained sections were imaged using a microscope image system (Zeiss, Axioscope 5). The stained areas were analyzed and counted by ImageJ software (Maryland, USA).

### Western Blots (WB) analysis

Cells or ground tissues were lysed in Ripa lysis solution (Beyotime, P0013C) for 1 h. The lysate was collected and the supernatant was collected by centrifugation at 4°C (12,000g, 25 min). The protein concentration in the supernatant was measured according to the BCA protein assay kit protocol (New Cell & Molecular Biotech, WB6501), and protein loading buffer (New Cell & Molecular Biotech, WB2001) was added to one-quarter volume of the supernatant and boiled for 10 min to denature the protein. Configure the electrophoresis gel according to the 10% ExpressCast PAGE Color Gel Express Kit instructions (New Cell & Molecular Biotech, P2012), add 3uL prestained color protein marker (New Cell & Molecular Biotech, P9005) or 30 mg of sample protein per well, and used the electrophoresis apparatus (LIUYI, DYCZ-40G) at a constant voltage of 90V for 2 hours. The separated proteins in the gel were transferred to PDVF membrane by transferring them through a transfer instrument (LIUYI, DYY-7C) at a constant current of 300mV for 2 hours. The gels were blocked for 1h in 2.5% milk in TBST and incubated for 16h at 4°C using diluted primary antibody. Table EV 2 showed the primary antibody information. 3 washes on the tilting shaker using TBST for 20 min each. Incubate with HRP-labeled goat anti-mouse IgG (Proteintech, PR30011) at 37°C for 1 h. Wash 3 times with TBST for 20 min each. The PDVF membranes were imaged by an imaging system (Bio-rad) according to the Enhanced Chemiluminescent (ECL; New Cell & Molecular Biotech, P2300) ptotocols.

### Lectin staining of Tissue sections

Paraffin sections were dehydrated, antigen-repaired, and blocked according to the experimental procedures in IHC, and then incubated overnight at 4°C using Bandeiraea Simplicifolia Lectin II (BSL-II; Vectorlabs, B-1215-2) diluted in PBS (dilution ratio 1:300) and washed three times with PBS for 3 min each. Prepare SABC working solution according to the instructions of the SABC-HRP kit (Beyotime, P0603) and was dropped on slides and incubated for 1h at room temperature. Prepare DAB working solution according to the instructions of DAB Staining Kit (ZSGB-BIO, ZLI-9018), and was dropped on slides and incubate for 10 min at room temperature, and terminate the incubation by washing twice with water. Dehydration and sealing were performed according to the experimental protocols in IHC. Imaging was performed using a microscope image system (Zeiss, Axioscope 5). The staining in the IHC images was analyzed and scored by the IHC Profiler plug-in in ImageJ software (Maryland, USA). Images were scored according to staining intensity versus staining area, High positive (3+), Positive (2+), Low Positive (1+) and Negative (0).

### Cell culture

Human colorectal cancer cell lines LOVO, NCM460, FHC, HT29, CACO2, RKO, HCT116, HCT8, SW480 were provided by NHC Key Laboratory of Carcinogenesis in Cancer Research Institute of Central South University. The cultures were incubated in DMEM medium (Biological Industries) with 10% fetal bovine serum (Gibco), 100 U/ml penicillin and 100 mg/ml streptomycin at 37°C and 5% CO2. LOVO cells were treated with 40uM Rosiglitazone (Meilunbio, MB1211, CAS: 122320-73-4), or 20uM GW9662 (MCE, HY-16578, CAS: 22978-25-2), or 2.5uM Kenpaullone (Selleck, S7919, CAS: 142273-20-9) for 24 hours.

### Plasmids, siRNA and Transfection

The siRNA and negative control (siControl) of MUC2, PPARg, KLF4 were purchased from RIBOBIO, where the MUC2-siRNA sequence was designed by RIBOBIO and the KLF4-siRNA sequence (UGAGAUGGGAACUCUUUGUGUAGGU) was obtained from Kazutoshi et al^15^. The plasmids pcDNA3.1-3xFlag-PPARg (NM_015869), pcDNA3.1-3xFlag-KLF4 (NM_001314052), pcDNA3.1-3xFlag-B3GNT6 (NM_138706), and the control blank plasmid pcDNA3.1-3xFlag were purchased from Youbio. LOVO cells were inoculated in six-well plates and incubated for 12 h. When the cell fusion rate reached 30-40%, the cells were prepared for transfection of siRNA when the cell fusion rate reached more than 50%. Transfection was performed according to the JetPRIME (Polyplus) procedure. After 12 hours of culture, the cytosol was changed and the cells were lysed after 48 hours for protein extraction or RNA purification.

### RNA isolation and Real-time quantitative PCR

The cells in the 6-well plate were washed with PBS and 1 ml of Trizol (Beyotime, R0016) was added to each well and the cells were collected after 5-10 min of lysis. Add 200μl chloroform, centrifuge at 4°C (15 min, 12000g), take the upper layer of liquid and add isopropyl alcohol and mix well, let stand at room temperature for 10 min, centrifuge at 4°C (10 min, 12000g). 75% ethanol wash the precipitate twice, centrifuge at 4°C (7 min, 7500g). Add 30-100 μl of enzyme-free water to dissolve the precipitate and detect the RNA concentration using a Nano Drop quantifier. Recerse transcription was performed the HiScript II qRT SuperMix for qPCR kit (Vazyme, R223-01). Real-time fluorescent quantitative PCR assays were performed using the SYBR qPCR Master Mix kit (Vazyme, Q321-02) with primers, the sequences of which are shown in Table EV 3.

### Cell viability assay

LOVO cells were seeded into 96-well plates with 8000 cells per well and incubated for 12 hours. Cell Counting Kit-8 (CCK8; New Cell & Molecular Biotech, C6005) was added after 0 h, 24 h, 48 h, 72 h, 96 h, and 120 h, respectively, and the absorbance at 450nm wavelength was measured by a microplate reader after 2 h of incubation at 37°C to analyze the change in cell proliferation Changes in cell proliferation capacity. Cell proliferation curves were obtained from at least five independent experiments.

### Clonogenic assay

LOVO cells were seeded into 6-well plates with 3,000 cells per well, and after 14 days of culture the cells were washed using PBS, fixed in 4% paraformaldehyde for 30 min, soaked in crystalline violet staining solution (Beyotime, C0121) for 10 min, washed in PBS and air-dried, and scanned for images. Clone formation results were obtained from at least 3 independent experiments.

### Transcription Factor Analysis

UCSC Genome Browser (http://genome.ucsc.edu) was used to obtain target gene’s promoter sequences (2000bp upstream of the transcription start site and 100bp downstream of the transcription start site). JASPAR (https://jaspar.genereg.net) was used to obtain the motifs of TFs and to align the motif of TFs with the promoter sequence, which is scanned to check whether the motif could be enriched. The ChIP-seq data and ChIP peak were analyzed on the Cistrome Data Browser (http://cistrome.org/db/) to predict to predict the binding sites of transcription factors to the promoter sequence of the target genes.

### Chromatin Immunoprecipitation (ChIP) Assay

LOVO cells were incubated for 10 min at 37°C in medium containing 1% formaldehyde and incubated with glycine for 10 min at room temperature to terminate protein cross-linking. Cell suspensions were prepared by adding PBS containing 1 mM PMSF. Cells were sonicated according to the protocols of the Chromatin Immunoprecipitation (ChIP) Assay Kit (Beyotime, P2078) and DNA fragments bound to transcription factors were obtained using antibodies to transcription factors as described in Table EV 2. proteins were digested according to the instructions and DNA fragments were purified using a DNA extraction kit (Vazyme, DC301-01). The predicted transcription factor binding fragments were amplified using a DNA polymerase kit (Vazyme, P505-D1) and the primer sequences are shown in Table EV 4. The amplified products were verified using qPCR or agarose gel electrophoresis.

### Luciferase reporter assay

Whole cell genomic DNA was extracted from LOVO cells using a genomic DNA kit (Tiangen, DP304) and sequences from 2 kb upstream to 100 bp downstream of the gene start codon were amplified using a DNA polymerase kit. Amplified fragments were cloned into pGL3-Basic plasmid. The transcription factor binding site was mutated using a mutagenesis kit (Vazyme, C215-01). The pGL3-Basic plasmid or mutant plasmid containing the gene promoter sequence was co-transfected into LOVO cells with the pRL-TK plasmid, and the cells were incubated for 2 days and then assayed for luciferase activity using a dua-luciferase reporter assay system (UEBlandy, UE-F6075). The firefly luciferase activity was normalized using renilla luciferase activity.

### Co-immunoprecipitation (Co-IP)

After transfection of pcDNA3.1-3xFlag-B3GNT6 plasmid in LOVO cells for 48 h, cells were lysed in cell lysis buffer for Western and IP (Beyotime, P0013) for 1 h. The lysate was collected and the supernatant was collected by centrifugation (12,000g, 25 min) at 4°C. Wash the anti-Flag immunomagnetic beads (Selleck, B26101) 3 times for 3 min each using cell lysis buffer for Western and IP on a DNA mixer. Mix anti-Flag immunomagnetic beads with 4ug IgG antibody (Proteintech, B900620) or 4ug B3GNT6 antibody (Cusabio, CSB-PA744392LA01HU) and incubate on the DNA Mixer for 3 hours. Add Cell Protein Lysate and incubate overnight at 4°C. Wash the beads 7 times for 5 min each using cell lysis buffer for Western and IP on the DNA Mixer. Discard the supernatant, add 20uL of cell lysis buffer for Western and IP with 5uL of Protein Loading Buffer (New Cell & Molecular Biotech, WB2001) and boil for 10 minutes to denature the proteins. Follow the subsequent Western Bot experiments.

### Lectin staining assay

After transfection of pcDNA3.1-3xFlag-B3GNT6 plasmid in LOVO cells for 48 h, the cells were lysed in cell lysis buffer for Western and IP for 1 h. The lysate was collected and centrifuged at 4°C (12,000g, 25 min) and the supernatant was collected. Wash Protein A/G immunoprecipitated magnetic beads (Selleck, B23201) 3 times for 3 min each using cell lysis buffer for Western and IP on a DNA mixer. Anti-Flag immunomagnetic beads were mixed with 4ug IgG antibody (Proteintech, B900620) or 4ug MUC2 antibody (Abclonal, A4767) and incubated on the DNA mixer for 3 hours. Add cellular protein lysate and incubate overnight at 4°C. Wash the beads 7 times for 5 minutes each using cell lysis buffer for Western and IP on the DNA Mixer. Discard the supernatant, add 20uL of cell lysis buffer for Western and IP with 5uL of Protein Loading Buffer and boil for 10 minutes to denature the proteins. After protein lysis, electrophoresis, transfer, and blocking according to the experimental steps in Western Blot, proteins were incubated overnight at 4°C using Bandeiraea Simplicifolia Lectin II (BSL-II; Vectorlabs, B-1215-2) diluted in PBS (dilution ratio 1:300) and washed three times for 3 min each in TBST. Configure SABC working solution according to the instructions of the SABC-HRP kit and incubate dropwise onto PDVF membranes for 1 h at room temperature. Prepare DAB working solution according to the instructions of DAB staining kit, and was dropped on PDVF membrane and incubate for 10 min at room temperature, and terminate the incubation by washing twice with TBST.

### StcE digestion assay

1 mL of medium with 5ug of StcE (Cusabio, CSB-EP530572EOD) was added per 1 million cells and incubated for 2 h at 37°C. The cells were incubated on ice according to Yao et al^16^ using 50 mM Tris-Cl (pH 8.0), 1% NP-40, 0.3% Triton X-10, 3% glycerol and 150 mM NaCl for 30 min and centrifuged at 4°C (25 min, 25,000 g) to obtain cellular proteins degraded by StcE. Followed by Western Blot experiments.

### Enzyme-linked immunosorbent assay (ELISA)

Collect LOVO cell supernatant, centrifuge at 1000g for 20 min at 2-8°C, and discard the precipitate. The amount of MUC2 in the cell supernatant was detected according to the instructions of MUC2 ELISA kit (Elabscience, E-EL-H0632c), and each experiment was repeated 3 times. Use Curve Expert software to plot the ELISA standard curve and calculate the concentration of target protein in the sample.

### Mice

All animal models were constructed in accordance with the Regulations for the Administration of Affairs Concerning Experimental Animals established by State Scientific and Technological Commission of The People’s Republic of China and passed the animal ethical review by the Department of laboratory animals of Central South University (ID: CSU-2022-0684). The animal cohort was all female C57BL/6J, provided by the Department of laboratory animals of Central South University, and housed in the Experimental Animal Center of Central South University. Mice were housed in SPF conditions, with ambient temperature at (23± 3) °C and on an alternating 12 h light-dark cycle. Mice were given a regular chow and bedding was changed every 2 days, and regular feed and bedding were provided by the Department of laboratory animals of Central South University.

In vivo, diets containing rosiglitazone (10 mg/kg) or GW9662 (10 mg/kg) were used to feed C57BL/6 mice. Two weeks later, IHC assays of PPARg, KLF4, B3GNT6 and MUC2 and BSL-II lectin staining were detected in intestinal tissues. MUC2 secretion level was analyzed in intestinal mucus using ELISA analysis.

### AOM/DSS model

After 1 week of acclimatization, each group was given Azoxymethane (AOM; 12 mg/kg) (Abiowell, AWH1007a, CAS:25843-45-2) subcutaneously, followed by Disuccinimidyl suberate (DSS; 3%) (Meilunbio, MB0842, CAS: 68528-80-3) solution ad libitum for 7 days after 1 week, then given normal drinking water for 2 weeks. Every 1 week of DSS water and 2 weeks of normal drinking water were used as 1 cycle. Mice in the experimental group were fed Rosiglitazone (10 mg/kg) daily from 3 days before AOM injection. The mice in the blank group were not given drug treatment. Six mice were randomly executed on the last day of each DSS cycle (day 23, day 44, and day 65 after AOM injection). Mice were euthanized when they lost 20-25% of their body weight, or when the size of the solid tumor exceeded 10% of the animal’s body weight, or when the tumor area of the solid tumor exceeded 2000 mm^3^, or when the diameter of the solid tumor exceeded 20mm in any one dimension. After the mice were executed, the intestinal lumen was opened along the longitudinal axis, and the number and diameter of tumors were recorded. Feces were collected from the intestinal cavity, and the mice were tested for fecal occult blood according to the Fecal Occult Blood Kit (Leagene Biotechnology) assay and the Disease Activity Index (DAI) score was determined by assessing the degree of daily weight loss.

### Histological scoring

HE staining of paraffin sections was performed according to the hematoxylin-eosin kit (Solarbio, G1120) staining method, and two independent pathologists scored the extent and degree of inflammatory involvement and the degree of crypt damage in the sections, according to the colon pathology scoring method of Horino et al^17^, as described in Table EV 1.

## Acknowledgments

We would like to thank to the Cancer Research Institute of Central South University for providing human colorectal cancer cell lines. This study was supported by grants from the National Natural Science Foundation of China (81872473 to P.C and 81972773 to Z.L.), the Natural Science Foundation of Hunan Province (No. 2019JJ40161 to Z.L.), and the Project of Improving the Diagnosis and Treatment Capacity of Hepatobiliary Pancreas and Intestine diseases in Hunan Province (Xiangwei [2019] No.118). The experimental equipment is provided by the Cancer Research Institute of Central South University and the Department of laboratory animals of Central South University.

## Author Contributions

YAY performed the experiments; analyzed the data and wrote the manuscript. YSY was responsible for animal feeding and mouse model construction. WX and YK performed bioinformatics analyses. JWC participated in collection and assembly of data. YFZ collected the intestinal tissue samples from patients. PGC and ZL were in charge of experiment design and manuscript review. All authors contributed to writing the manuscript and final approval of the manuscript.

## Data availability statement

Data are available on reasonable request. All data are available from the corresponding author on request.

## Expanded View Figure legends

**Figure EV 1.**
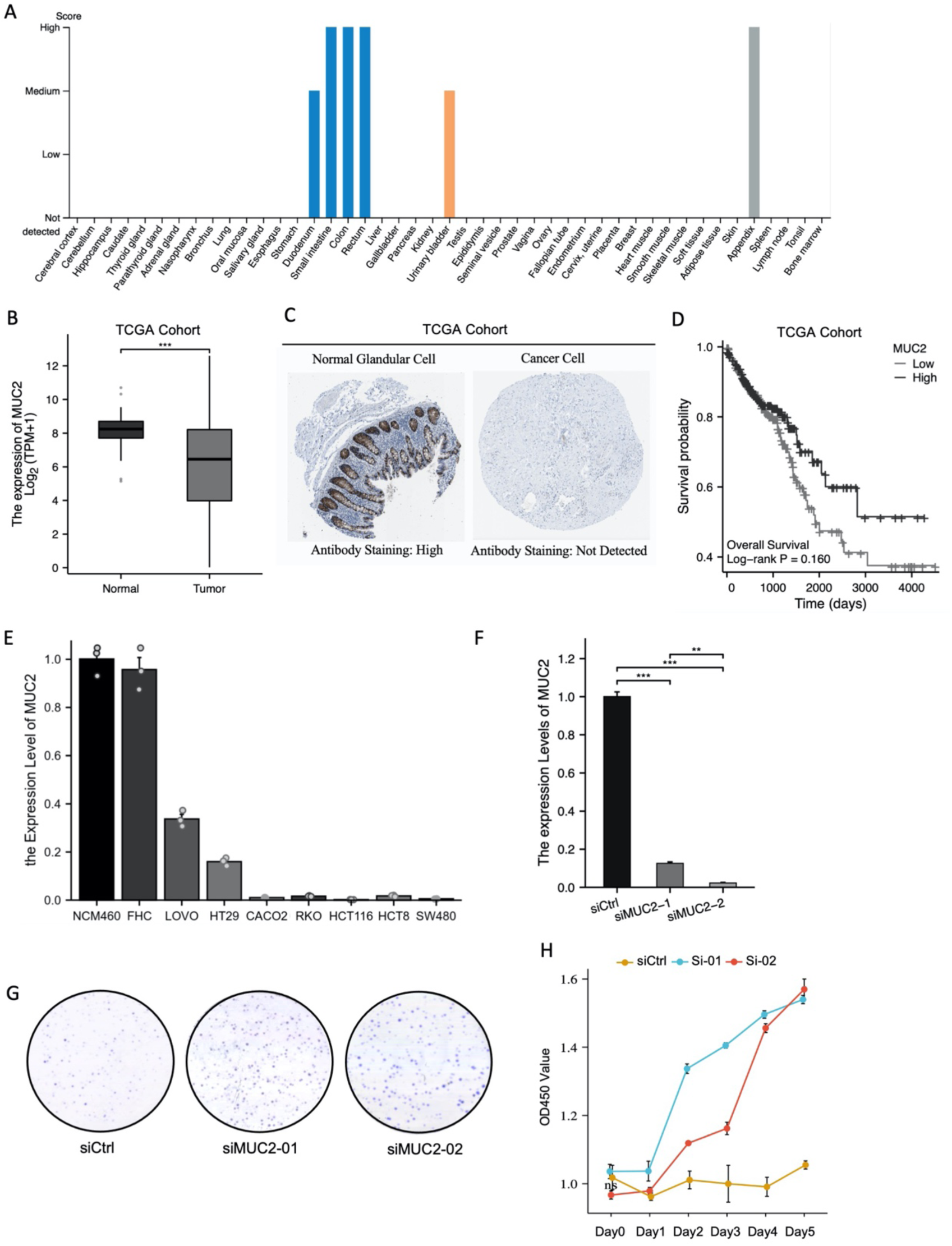
MUC2 Knock-down increases the proliferation ability of colorectal cancer cell. (A) Expression and distribution of MUC2 in different organs of human body.. (B) Boxplot analysis of MUC2 expression level in tumors and normal tissues in CRC patients in TCGA. (C) IHC staining of MUC2 expression of CRC patients in HumanProteinAtlas database. (D) Survival analysis for MUC2-High and MUC2-Low patients with CRC in TCGA.(E) QPCR detection of the relative abundance of the expression level of MUC2 in different colorectal cancer cell lines. (F) QPCR detection of the relative abundance of the expression level of MUC2 after siRNA transfection into LOVO cells (n=3, Ctrl; n=3, siMUC2-1; n=3, siMUC2-2). (G) Colony formation of LOVO cells following transfection with siRNA (n=3, Ctrl; n=3, siMUC2-01; n=3, siMUC2-02). (H) Cell proliferation of LOVO cells following transfection with siRNA was evaluated by CCK8 (n=5, Ctrl; n=5, siMUC2-01; n=5, siMUC2-02). Data were analyzed by ordinary one-way ANOVA with Tukey’s multiple comparisons. The error bars indicate means ±standard error (SEM). *p<0.05, **p<0.01, ***p<0.001. ANOVA, analysis of variance; IHC, immunohistochemistry; CRC, Colorectal Cancer, qPCR, quantitative polymerase chain reaction; H&E, hematoxylin-eosin; CCK8, cell counting kit-8.

**Figure EV 2.**
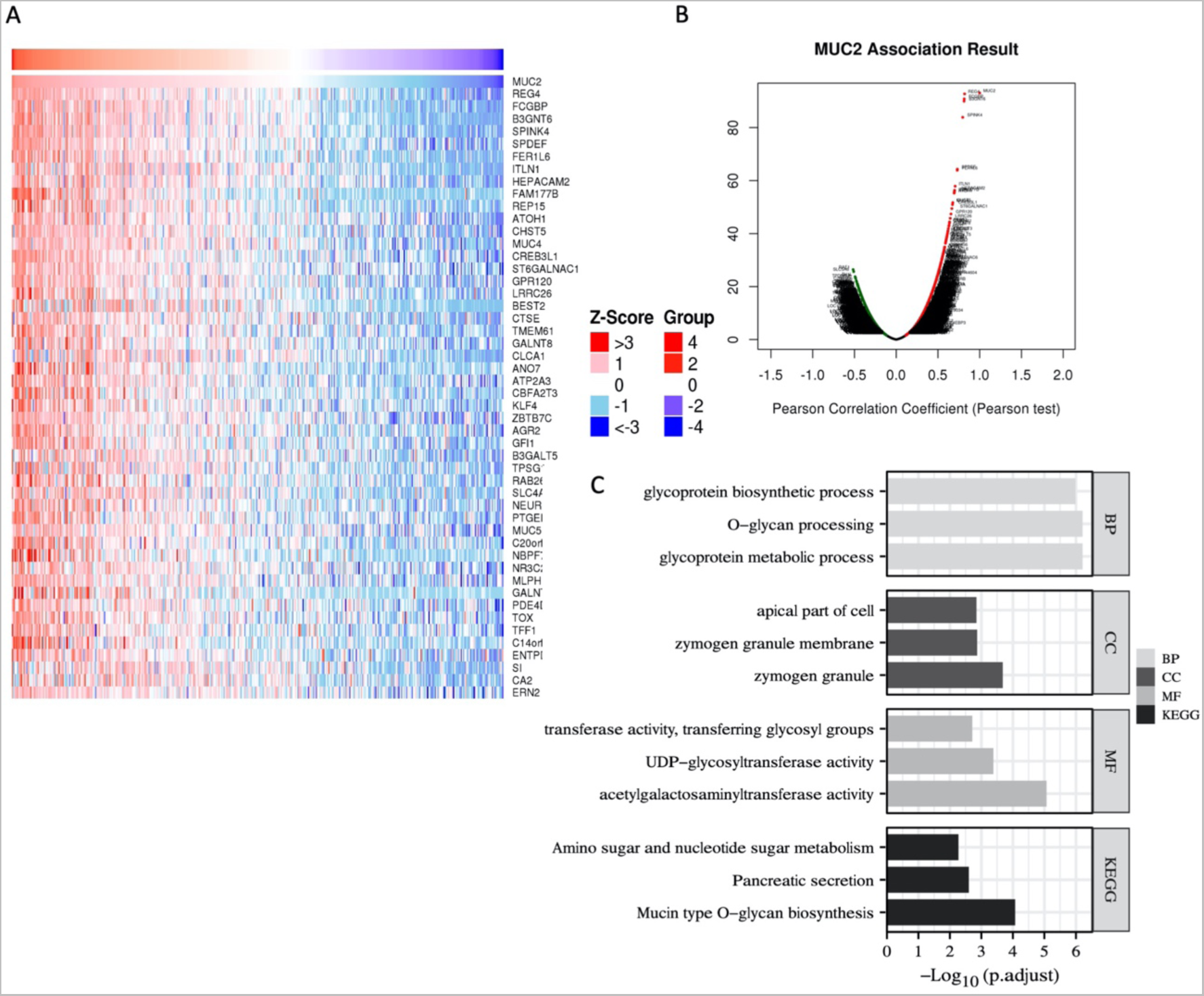
B3GNT6 was the key molecule in MUC2 O-glycosylation pathway. (A-B) The Pearson correlation between MUC2 and other genes in TCGA dataset was visualized by heatmap and volcano plot. Red dots indicated significantly positive correlation with MUC2 and blue dots indicated significant negative correlation with MUC2. (C) Significantly enriched GO annotations and KEGG pathways of the genes related to MUC2 expression. GO, Gene Ontology; KEGG, Kyoto Encyclopedia of Genes and Genomes.

**Figure EV 3.**
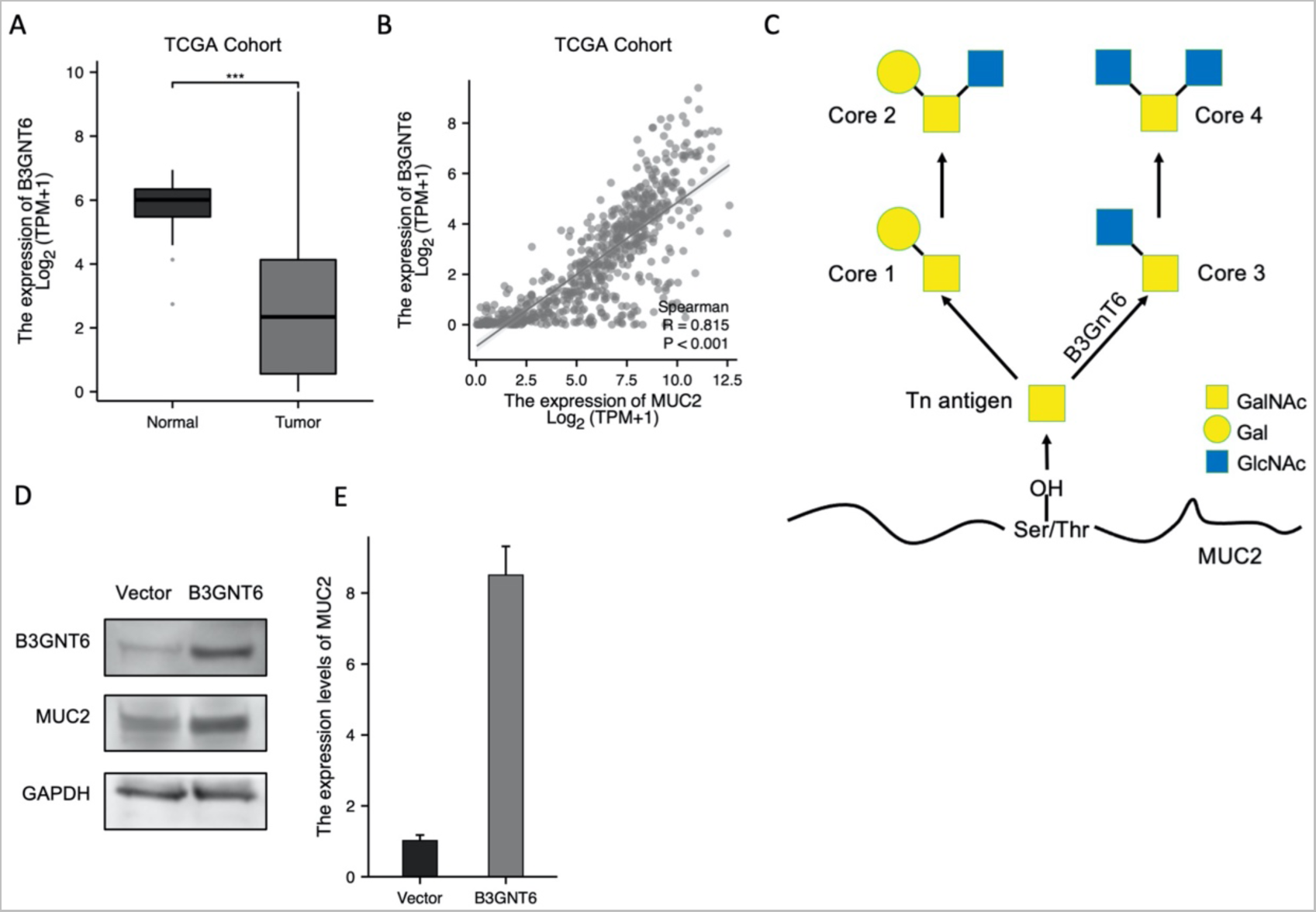
B3GNT6 enhances the expression of MUC2. (A) Boxplot analysis of B3GNT6 expression level in tumors and normal tissues in CRC patients in TCGA. (B) A scatter plot and Pearson correlation score showing a positive correlation between MUC2 and B3GNT6 in TCGA. (C) Schematic diagram of the synthetic pathway of O-glycosylation. Data were analyzed by ordinary one-way ANOVA with Tukey’s multiple comparisons. (D-E) QPCR and Western Blot assay to evaluate the expression level of MUC2 after pcDNA3.1-B3GNT6 plasmid transfection into LOVO cells (n=3, Ctrl; n=3, B3GNT6-OE) The error bars indicate means ±standard error (SEM). *p<0.05, **p<0.01, ***p<0.001. Data were analyzed by ordinary one-way ANOVA with Tukey’s multiple comparisons. The error bars indicate means ±standard error (SEM). *p<0.05, **p<0.01, ***p<0.001. ANOVA, analysis of variance; qPCR, quantitative polymerase chain reaction.

**Figure EV 4.**
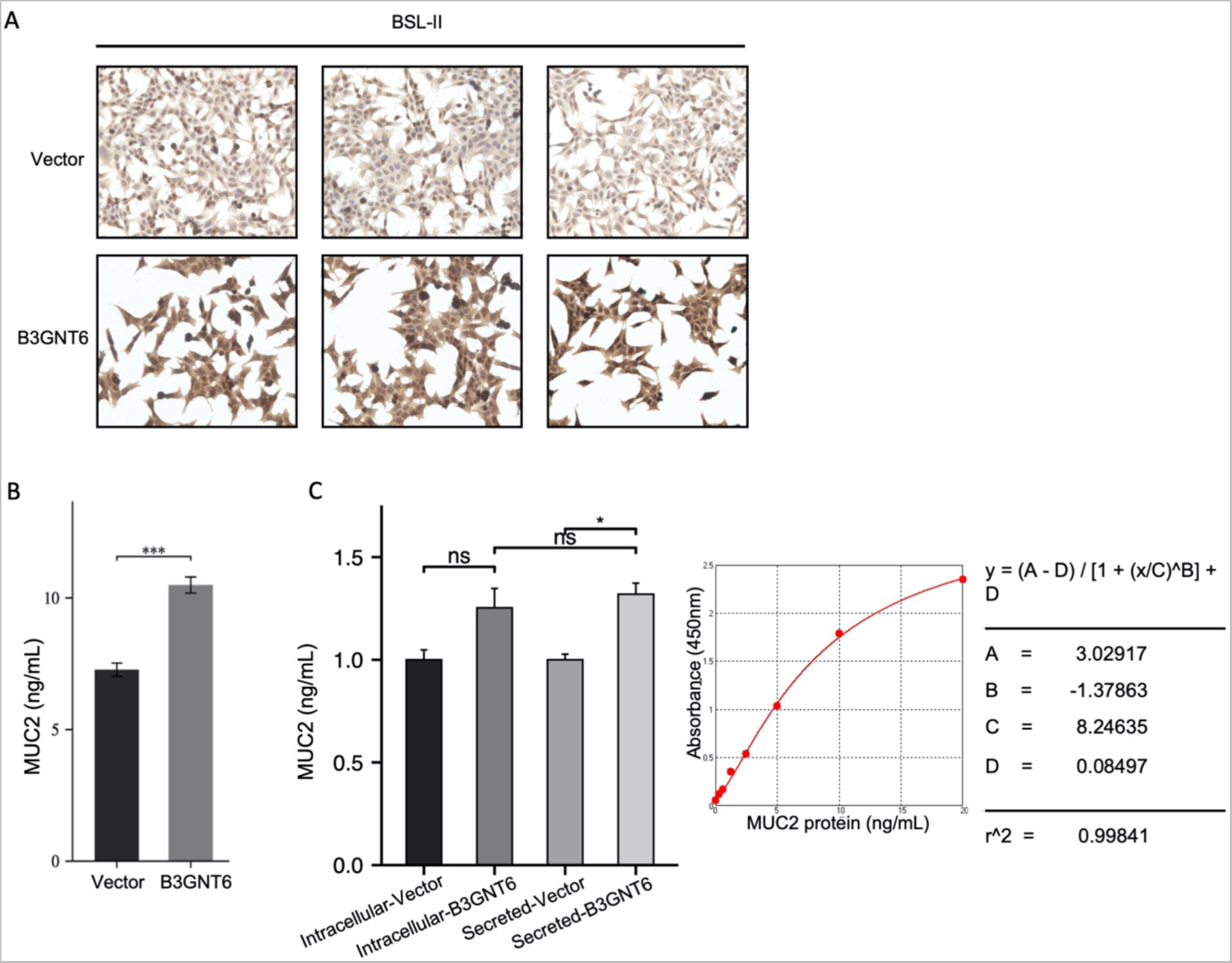
B3GNT6 increases the level of MUC2 secretion and Core3 modification. (A) Lectin staining using BSL-II and in LOVO cells after transfection of pcDNA3.1-B3GNT6 plasmid into LOVO cells (n=3, Ctrl; n=3, B3GNT6-OE). Scale bars, 50 µm. (B) MUC2 secretion was measured by ELISA after transfection of pcDNA3.1-B3GNT6 plasmid into LOVO cells (n=5, Ctrl; n=5, B3GNT6-OE). (C) ELSIA assay evaluated the level of MUC2 in cell lysate and cell supernatant after pcDNA3.1-B3GNT6 plasmid transfection into LOVO cells (n=5, Intracellular-Ctrl; n=5, Intracellular-B3GNT6; n-5, Secreted-Ctrl; n=5, Secreted-B3GNT6). Data were analyzed by ordinary one-way ANOVA with Tukey’s multiple comparisons. The error bars indicate means ±standard error (SEM). *p<0.05, **p<0.01, ***p<0.001. ANOVA, analysis of variance; OE, Overexpression; ELISA, enzyme-linked immunosorbent assay.

**Figure EV 5.**
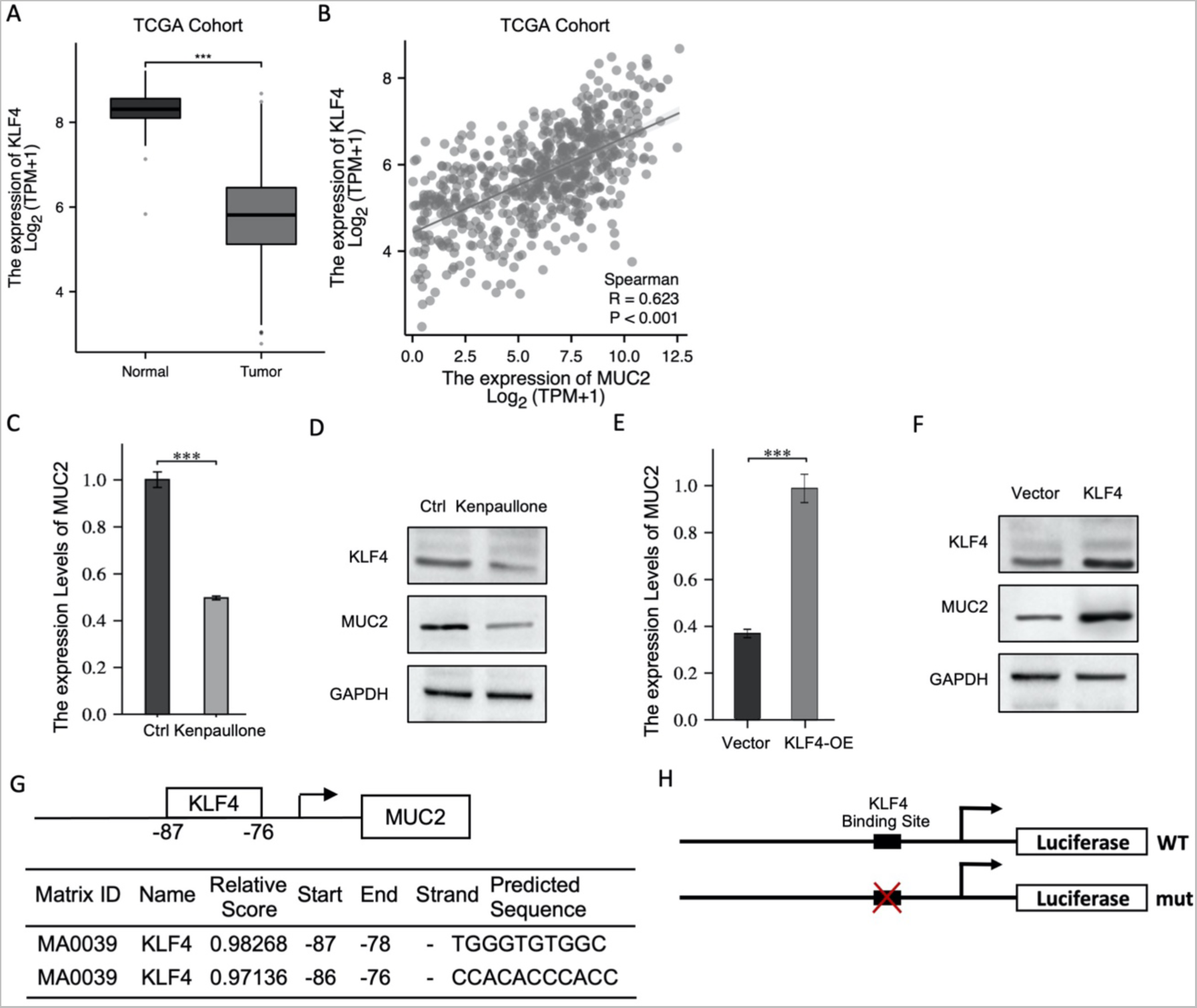
KLF4 is involved in the regulation of MUC2 transcription. (A) Boxplot analysis of KLF4 expression level in tumors and normal tissues in CRC patients in TCGA. (B) A scatter plot and Pearson correlation score showing a positive correlation between MUC2 and KLF4 in TCGA. (C) QPCR and (D)Western Blot assay to evaluate the expression level of MUC2 in LOVO cells after administration of 2.5uM Kenpaullone (n=3, Ctrl; n=3, Kenpaullone). (E) QPCR and (F) Western Blot assay to evaluate the expression level of MUC2 after pcDNA3.1-KLF4 plasmid transfection into LOVO cells (n=3, Ctrl; n=3, KLF4-OE). (G) Schematic diagram of KLF4 binding site in the MUC2 promoter region. (H) Schematic diagram of MUC2 promoter reporter plasmids. PGL3basic-MUC2 promoter-WT plasmid was constructed by inserting MUC2 promoter sequence into pGLBasic luciferase reporter plasmid, and the mutation site (pGL3basic-MUC2 promoter-mut) was constructed by homologous recombination. Data were analyzed by ordinary one-way ANOVA with Tukey’s multiple comparisons. The error bars indicate means ±standard error (SEM). *p<0.05, **p<0.01, ***p<0.001. ANOVA, analysis of variance, CRC, Colorectal Cancer.

**Figure EV 6.**
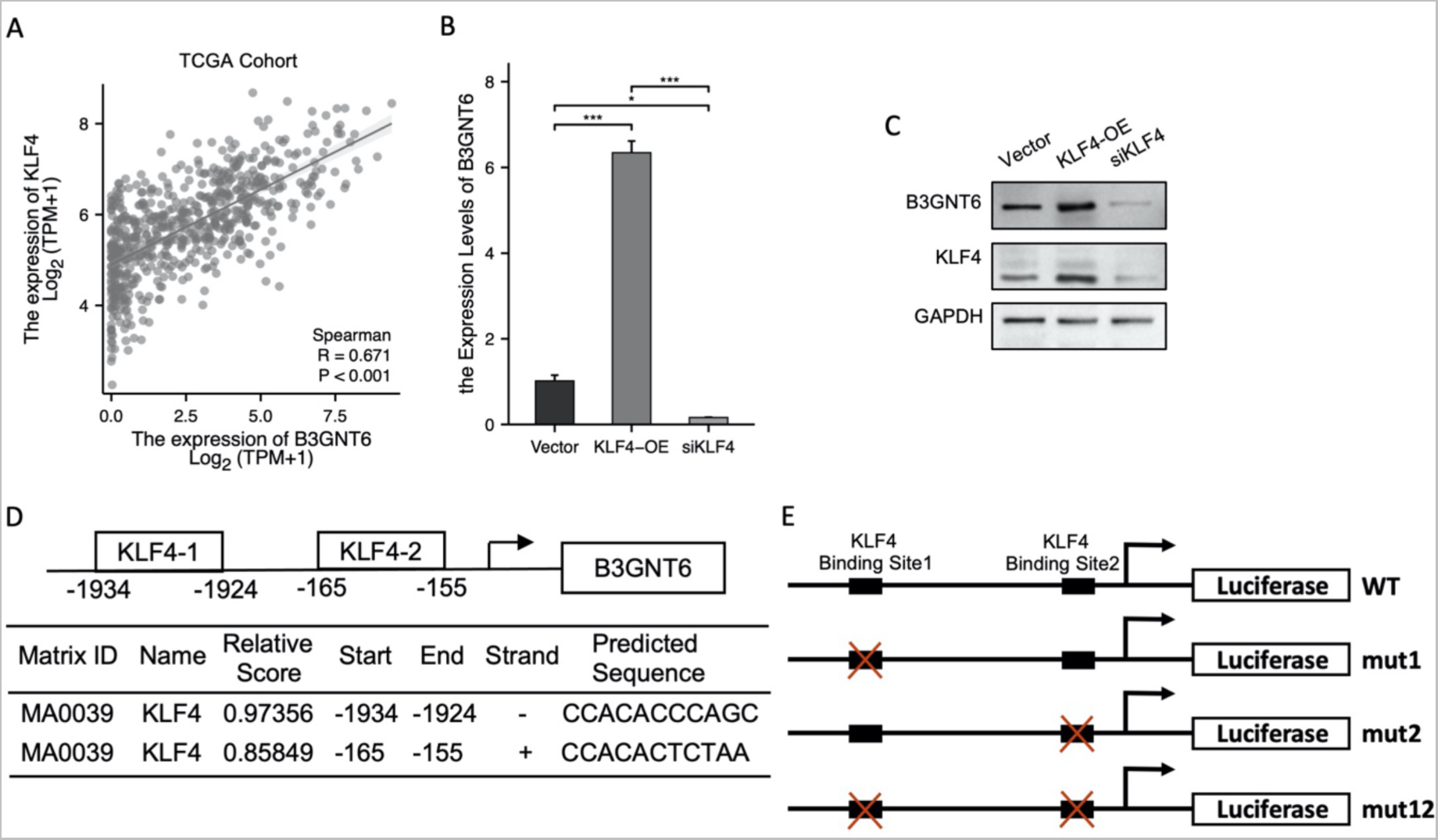
Prediction of binding sites for KLF4 binding MUC2 and B3GNT6 promoter region. (A) A scatter plot and Pearson correlation score showing a positive correlation between B3GNT6 and KLF4 in TCGA. (B-C) QPCR and Western Blot to detect the expression of B3GNT6 after transfection pcDNA3.1-KLF4 plasmid or siRNA-KLF4 into LOVO cells (n=3, Ctrl; n=3, KLF4-OE; n=3, siRNA-KLF4). (D) Schematic diagram of KLF4 binding site in the B3GNT6 promoter region. (E) Schematic diagram of B3GNT6 promoter reporter plasmids. PGL3basic-B3GNT6 promoter-WT plasmid was constructed by inserting B3GNT6 promoter sequence into pGLBasic luciferase reporter plasmid. pGL3basic-Mutated plasmids were constructed by mutating two binding sites separately or at the same time (PGL3basic-B3GNT6 promoter-mut1: 1924-1934bp; PGL3basic-B3GNT6 promoter-mut2: 155-165bp). Data were analyzed by ordinary one-way ANOVA with Tukey’s multiple comparisons. The error bars indicate means ±standard error (SEM). *p<0.05, **p<0.01, ***p<0.001. ANOVA, analysis of variance; qPCR, quantitative polymerase chain reaction.

**Figure EV 7.**
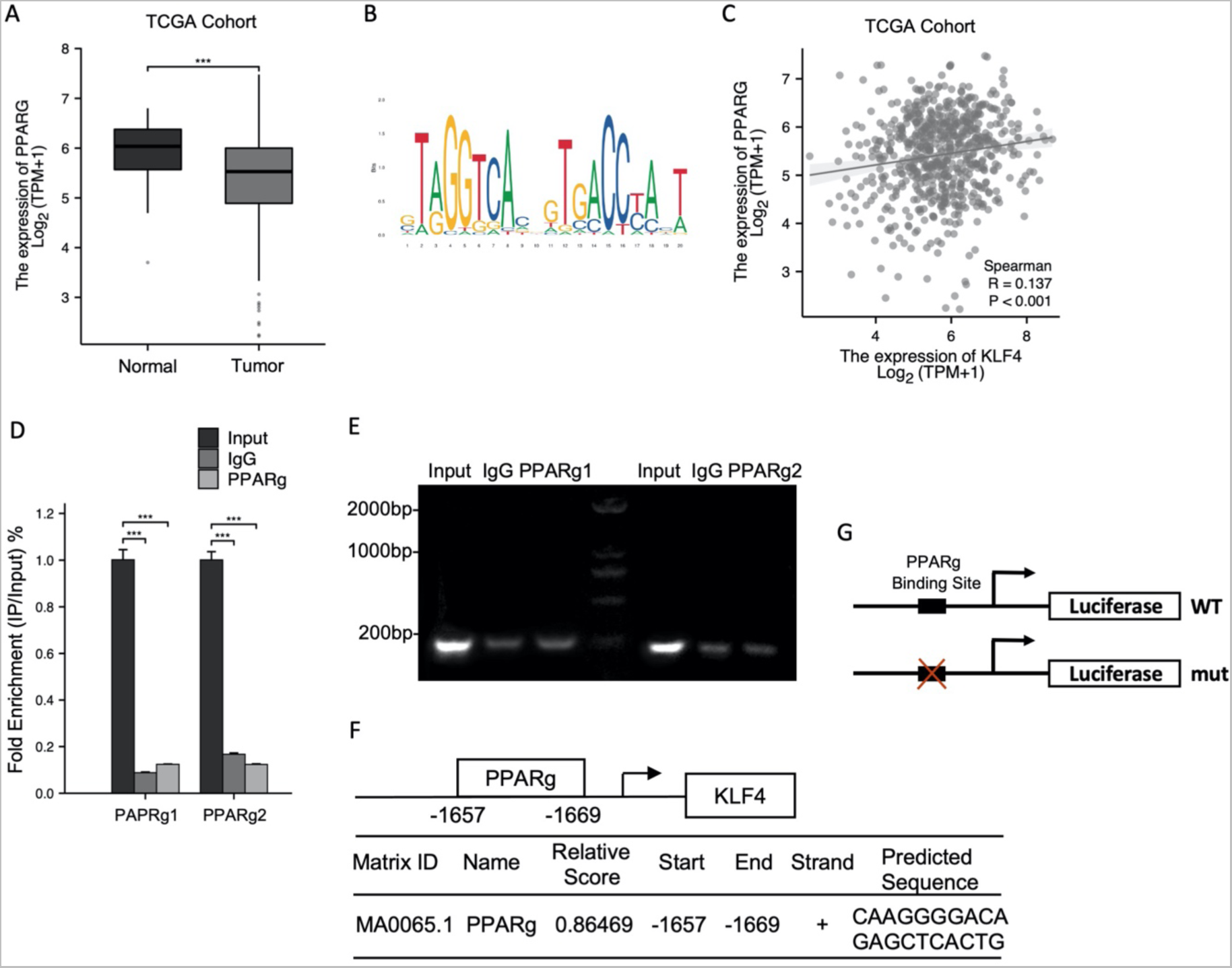
Prediction of binding sites for PPARg binding MUC2 and KLF4 promoter region. (A) Boxplot analysis of PPARg expression level in tumors and normal tissues in CRC patients in TCGA. (B) Sequence logo of PPARg binding sites based on the Jaspar database. (C) A scatter plot and Pearson correlation score showing a positive correlation between PAPRg and KLF4 in TCGA. (D-E) ChIP enrichment was quantified using PCR and qPCR analysis to detect PPARg-binding sites (n=3, Input; n=3, IgG; n=3, CHIP-PPARg). Binding site1 and site2 were 1959-1969bp and 517-535bp upstream of ATG initiation codon respectively. (F) Schematic diagram of PPARg binding site in the MUC2 promoter region. (G) Schematic diagram of KLF4 promoter reporter plasmids. PGL3basic-KLF4 promoter-WT plasmid was constructed by inserting KLF4 promoter sequence into pGLBasic luciferase reporter plasmid, and the mutation site (pGL3basic-KLF4 promoter-mut) was constructed by homologous recombination. Data were analyzed by ordinary one-way ANOVA with Tukey’s multiple comparisons. The error bars indicate means ±standard error (SEM). *p<0.05, **p<0.01, ***p<0.001. ANOVA, analysis of variance; ChIP, chromatin immunoprecipitation; qPCR, quantitative polymerase chain reaction.

**Figure EV 8.**
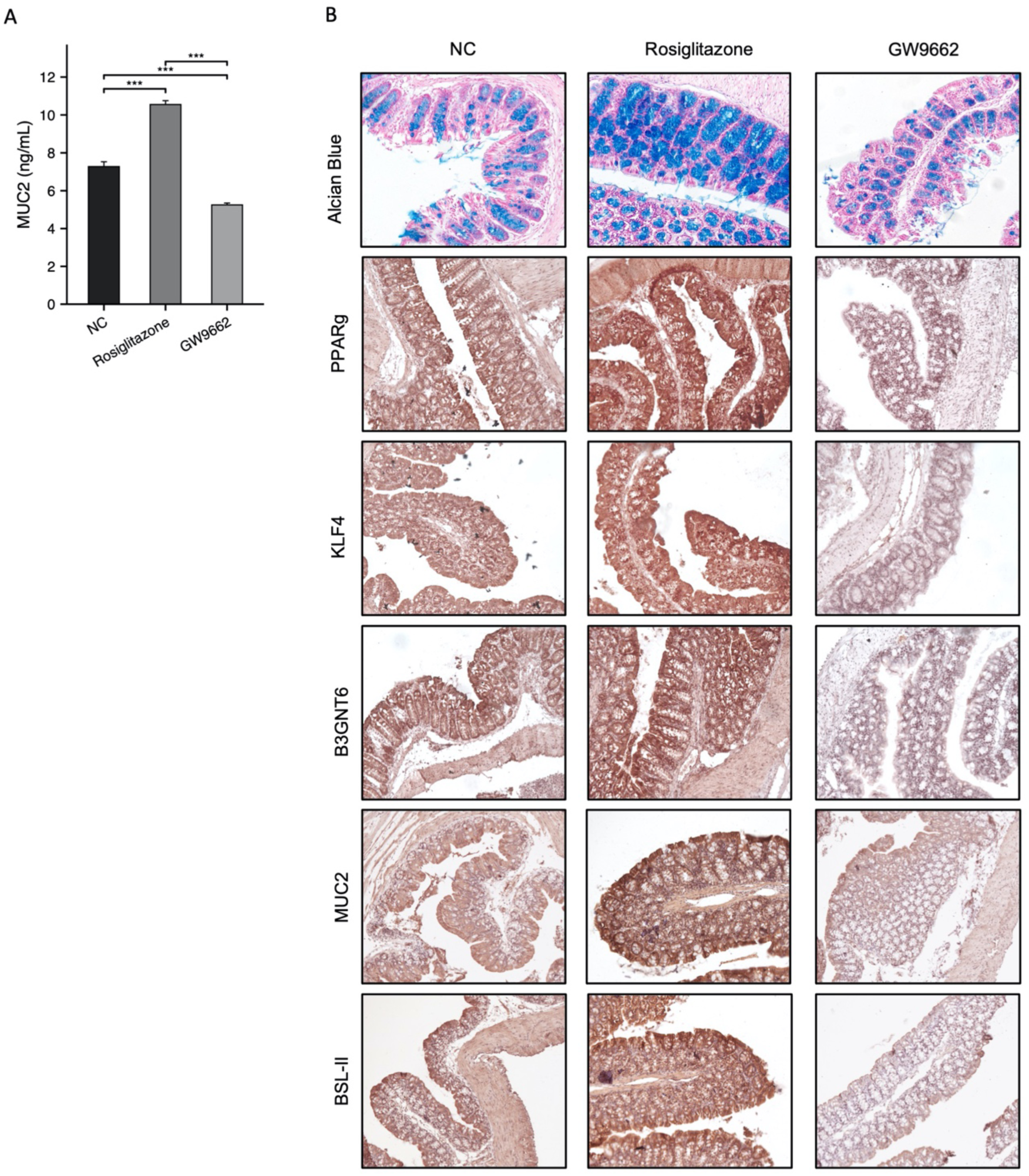
Effects of PPARg agonist rosiglitazone and inhibitor GW9662 on the expression of downstream molecules. (A) ELSIA assay evaluated the level of MUC2 in cell supernatant after treated with Rosiglitazone (40uM) or GW9662 (20uM) (n=5, Ctrl; n-5, Rosiglitazone; n=5, GW9662). (B) H&E, IHC staining of PPARg, KLF4, B3GNT6, MUC2 and BSL-II lectin staining of Core3 in the C57BL/6 treated with Rosiglitazone (10mg/kg) or GW9662 (10mg/kg) (n=3, Ctrl; n=3, Rosiglitazone; n=3, GW9662). Scale bars, 50 µm. Data were analyzed by ordinary one-way ANOVA with Tukey’s multiple comparisons. The error bars indicate means ±standard error (SEM). *p<0.05, **p<0.01, ***p<0.001. ANOVA, analysis of variance; H&E, hematoxylin-eosin; IHC, immunohistochemistry; CRC, Colorectal Cancer.

**Figure EV 9.**
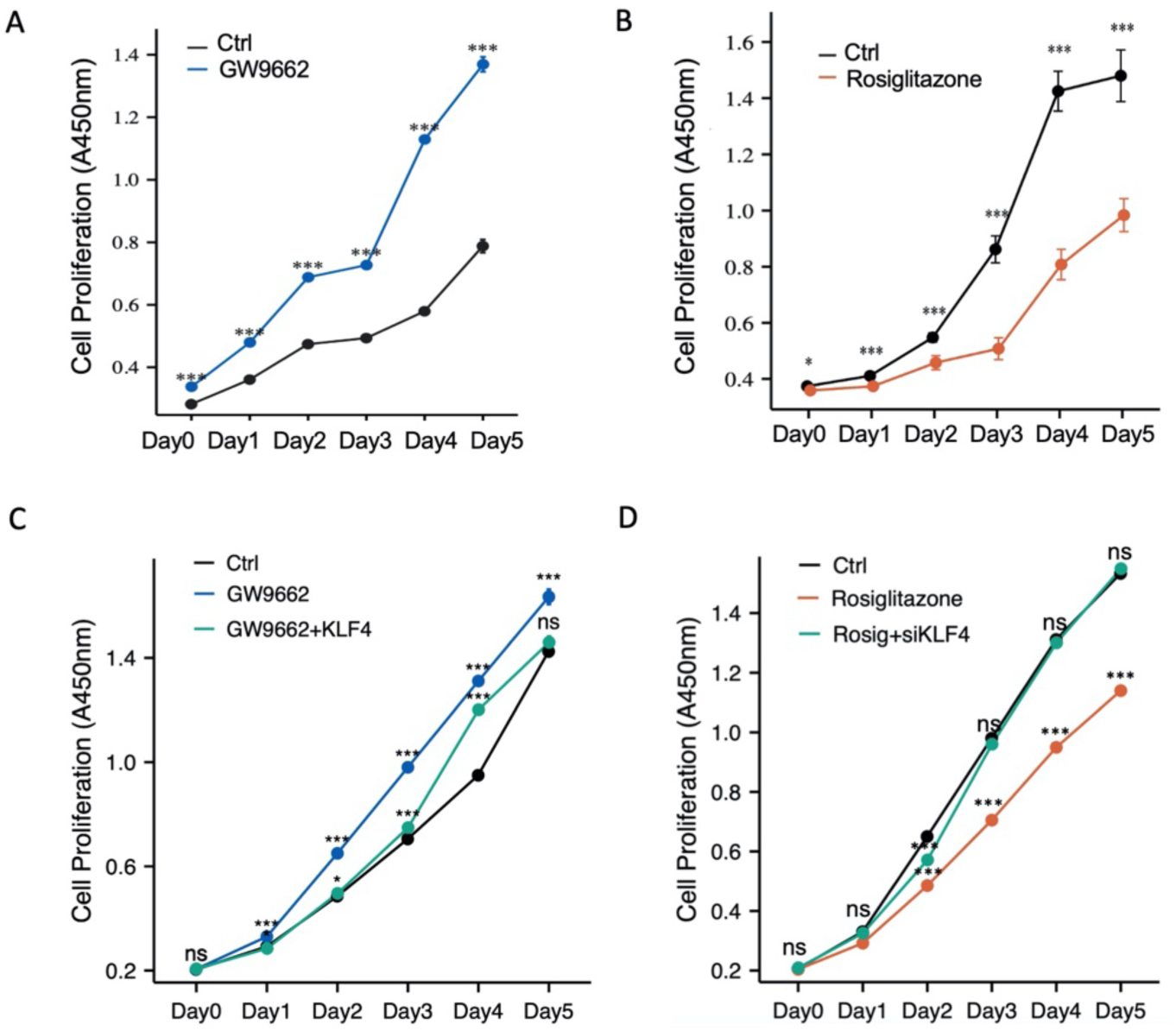
Rosiglitazone can inhibit the proliferation of colorectal cancer cells. (A-B) Cell proliferation of LOVO cells following treated with GW9662 (20uM) or Rosiglitazone (40uM) was evaluated by CCK8. (C-D) Cell proliferation of LOVO cells was evaluated by CCK8 after treated with GW9662 (20uM) and pcDNA3.1-KLF4 plasmid transfection, or Rosiglitazone (40uM) and siRNA-KLF4 transfection. Data were analyzed by ordinary one-way ANOVA with Tukey’s multiple comparisons. The error bars indicate means ±standard error (SEM). *p<0.05, **p<0.01, ***p<0.001. ANOVA, analysis of variance; CCK8, cell counting kit-8.

